# Generation of hiPSC-derived brain microvascular endothelial cells using a combination of directed differentiation and transcriptional reprogramming strategies

**DOI:** 10.1101/2024.04.03.588012

**Authors:** Aomeng Cui, Ronak Patel, Patrick Bosco, Ugur Akcan, Emily Richters, Paula Barrilero Delgado, Dritan Agalliu, Andrew A. Sproul

## Abstract

The blood-brain barrier (BBB), formed by specialized brain microvascular endothelial cells (BMECs), regulates brain function in health and disease. *In vitro* modeling of the human BBB is limited by the lack of robust hiPSC protocols to generate BMECs. Here, we report generation, transcriptomic and functional characterization of reprogrammed BMECs (rBMECs) by combining hiPSC differentiation into BBB-primed endothelial cells and reprogramming with two BBB transcription factors FOXF2 and ZIC3. rBMECs express a subset of the BBB gene repertoire including tight junctions and transporters, exhibit stronger paracellular barrier properties, lower caveolar-mediated transcytosis, and similar p-Glycoprotein activity compared to primary HBMECs. They can acquire an inflammatory phenotype when treated with oligomeric Aβ42. rBMECs integrate with hiPSC-derived pericytes and astrocytes to form a 3D neurovascular system using the MIMETAS microfluidics platform. This novel 3D system resembles the *in vivo* BBB at structural and functional levels to enable investigation of pathogenic mechanisms of neurological diseases.

## INTRODUCTION

The blood-brain barrier (BBB), formed by brain microvascular endothelial cells (BMECs), is critical for the central nervous system (CNS) function. The BBB function is impaired in many neurological disorders but also it imposes a problem for delivery of therapeutics to the CNS^1^. Interactions of BMECs with pericytes and astrocytes via the extracellular matrix (ECM) are critical to assemble the neurovascular unit (NVU), and induce and maintain the BBB properties ranging from: (1) the presence of tight junctions (TJs) which restrict paracellular flux; (2) reduced transcytosis due to the presence of few caveolae; (3) specialized transporters that regulate the passage of nutrients and restrict harmful pathogens; to (4) low expression of leukocyte adhesion molecules (LAMs)^2–6^. Rodent models, although useful for studying mechanisms of CNS diseases, fail to capture patient heterogeneity inherent in multi-factorial neurological disorders and their NVU cells differ from those of humans at the molecular level^7–9^. For example, the mitigating effects of patient comorbidities on traumatic brain injury (TBI) are not captured in rodents; rodent models also do not recapitulate the genetic diversity of Alzheimer’s disease (AD) patients^10–12^. Additionally, rodents have greater P-glycoprotein drug efflux activity than humans, leading to an underestimation of drug toxicity^9,13^. There is a need to develop *in vitro* human NVU systems with robust BBB properties to model neurological diseases and develop safe, effective therapeutics capable of penetrating the CNS^14–17^.

Primary human BMECs (HBMECs) are scarce, heterogenous, and dedifferentiate in culture^18–20^. Human induced pluripotent stem cells (hiPSCs) are a promising alternative due to their reproducibility, scalability, and ability to model various pathological conditions. Over the past decade, several protocols have claimed to generate a homogenous population of “brain” endothelial cells (ECs), termed iPSC-derived BMECs (iBMECs), with high trans-endothelial electrical resistance (TEER) and expression of TJ proteins^21–27^. We recently demonstrated that iBMECs are transcriptomically and functionally analogous to epithelial cells^28^. iBMECs also fail to respond to inflammatory stimuli, limiting their ability to model neuroinflammatory or neurodegenerative diseases^29^. Recent methods have used extended passaging of endothelial progenitor cells (EPCs) to generate ECs (EPC-ECs), with good barrier properties but limited response to inflammatory stimuli^29,30^. Our current understanding of how the human BBB changes in pathological states is limited in part by our ability to generate *bona-fide* brain ECs.

Comparisons of transcriptome and chromatin accessibility of ECs from various organs have identified several transcription factors (TFs) essential for BMEC identity including FOXF2, FOXQ1, LEF1, and ZIC3^31,32^. FOXF2 is expressed by ECs^31,32^ and pericytes^33^, and is critical to maintain BBB tight junction integrity and low transcytosis^33^. ZIC3 is upregulated during development, suggesting a potential role in BBB establishment^32,34^. Overexpression of FOXF2 and ZIC3 in primary human umbilical vein endothelial cells (HUVECs) is sufficient to upregulate a limited set of BBB-specific genes^31^, so we hypothesized that reprogramming iPSC-derived ECs with FOXF2 and ZIC3 would enable the acquisition of robust BBB properties.

Here, we describe our novel method to generate iPSC-derived reprogrammed BMECs (rBMECs) by combining an iPSC differentiation step into BBB-primed ECs (bpECs)^35^ with FOXF2 and ZIC3-mediated transcriptional reprogramming. rBMECs have higher levels of key BBB transcripts and proteins compared to primary HBMECs and bpECs, exhibit higher TEER, lower transcytosis and equivalent efflux transport compared to HBMECs at the functional level, and respond to angiogenic stimuli. rBMECs respond robustly to oligomerized amyloid-β (oAβ42) by upregulating pro-inflammatory LAMs. Finally, rBMECs cocultured in a 3D microfluidic system with iPSC-derived astrocytes and pericytes form an NVU, demonstrate high TEER levels, and low barrier permeability to tracers, closely resembling the *in vivo* human NVU/BBB. Taken together, rBMECs enable faithful *in vitro* modeling of the human BBB for future studies of disease pathogenesis and development of putative clinical therapies.

## RESULTS

### Reprogramming iPSC-derived bpECs into rBMECs

During mammalian embryonic development, the mesodermal layer gives rise to ECs that form the primitive perineural plexus and the CNS vasculature, under the guidance of vascular endothelial growth factor A (VEGF-A) and Wnt/β-catenin signaling (reviewed in^36^). We hypothesized that iPSC-derived rBMECs could be generated using a two-step approach mirroring development (**Figure 1A**). In the first step, iPSCs were differentiated into what we term BBB-primed ECs (bpECs) by sequential exposure first to BMP4/bFGF to generate a mesodermal intermediate followed by VEGF-A/Thymosin β4 using a published protocol with minor modifications^35^ (**Figures 1A, S1A**). bpECs were purified from other mesodermal cells by CD31^+^/VE-cadherin^+^ expression using fluorescence activated cell sorting (FACS) on day 10 of differentiation (Sort A) and expanded on collagen IV/fibronectin-coated dishes in human endothelial serum-free media supplemented with 1% plasma-derived platelet poor serum, VEGF-A, bFGF and SB431542 (**Figures 1A, B, S1A**). In the second stage, bpECs were transduced with lentiviruses expressing FOXF2 and ZIC3. Transduced and reprogrammed bpECs were grown for 14 days and re-purified by FACS (Sort B) to generate a pure population of rBMECs that could be reliably expanded for five passages after re-purification (**Figures 1A, B, S1A**). By Sort B, bpECs expressed lower levels of CD31 and VE-CADHERIN (CDH5) compared to rBMECs (**Figure 1C**). Sort A efficiency was variable for various iPSC lines ranging from 1.3-5.1% for IMR90 (cl.4), 3.9-7.4% for FA0000010 (FA10), and 2.4-3.3% for NCRM5 **(Figure S1C)**; Sort B efficiency was also variable ranging from 56.6-96.1% for IMR90, 83.7-92.3% for FA10, and 34.9-42.6% for NCRM5 (**Figure S1D**).

**Figure 1.**
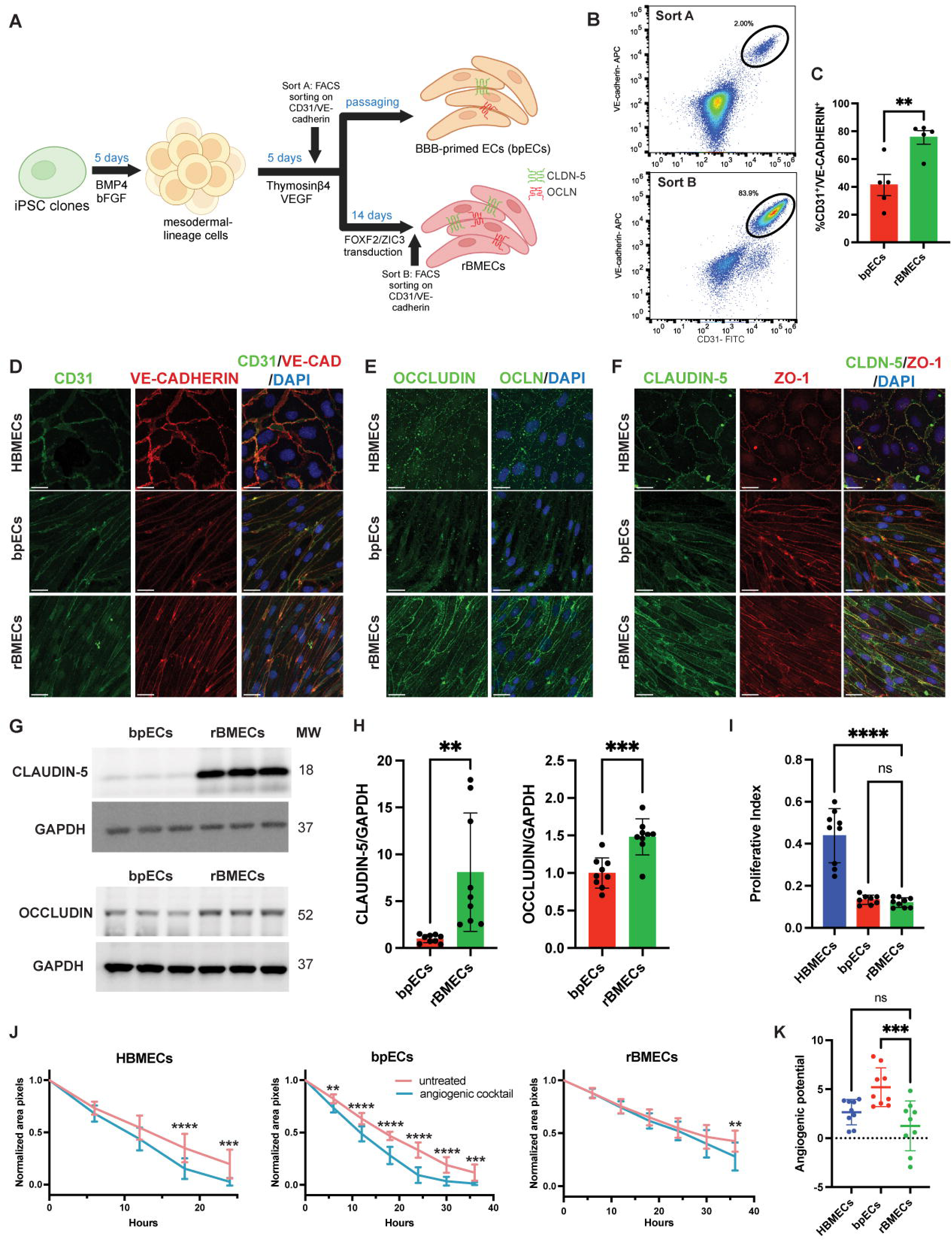
rBMECs generated through FOXF2/ZIC3 transcriptional reprogramming express key EC markers and respond to angiogenic stimuli. **A)** Schematic diagram for the generation of BBB primed ECs (bpECs) and reprogrammed BMECs (rBMECs) from human induced pluripotent stem cells (hiPSCs). **B)** Representative flow cytometry plots showing the fraction of CD31^+^/VE-CADHERIN^+^ ECs isolated after 10 days of the initial differentiation (sort A; bpECs) and 14 days after FOXF2/ZIC3 transduction (sort B; rBMECs). **C)** Dotted bar graph of the fraction of CD31^+^/VE-CADHERIN^+^ bpECs and rBMECs at 14 days post FOXF2/ZIC3 transduction; n = 5, unpaired t-test, **p<0.01. **D-F)** Representative immunofluorescence staining of HBMECs, bpECs and rBMECs for EC (CD31, VE-CADHERIN) and TJ (OCCLUDIN, CLAUDIN-5, ZO-1) proteins. DAPI (blue in merged images) labels the nuclei. Scale bar = 25 μm. **G-H)** Representative western blots and quantification for CLAUDIN-5 and OCCLUDIN expression. Mean ± SD, n = 9 from three experiments, unpaired t-test, **p<0.1, ***p<0.001. **I)** Proliferative index quantification after cell migration assay; n = 8-9 from three experiments, one-way ANOVA, ****p<0.0001. **J)** Quantification of the normalized area (pixels) of the wound region in the scratch assay; repeated measures ANOVA with Bonferroni correction; n = 9 from three experiments, *p<0.05, **p<0.1, ***p<0.001, ****p<0.0001. **K)** Quantification of angiogenic potential; n = 9 from three experiments, one-way ANOVA, ns p>0.05, ***p<0.001. (See also **Figure S1**).

### rBMECs express key endothelial and BBB markers and respond to angiogenic stimuli

To characterize rBMECs phenotypically and functionally, we first examined the expression of several EC and BBB proteins. The bpECs and rBMECs derived from all three iPSC lines expressed high levels of key EC identity proteins [e.g. VE-CADHERIN, platelet/endothelial cell adhesion molecule (PECAM1/CD31)] and BBB TJ proteins [CLAUDIN-5, OCCLUDIN, ZONULA OCCLUDENS-1 (ZO-1)] by immunofluorescence (IF) staining (**Figures 1D-F, S1E-J**). The rBMECs had higher CLAUDIN-5 (8.1 fold) and OCCLUDIN (1.5 fold) protein levels compared to bpECs by western blot (**Figure 1G-H**). Importantly, rBMECs did not express epithelial cellular adhesion molecule (EPCAM) as assessed by flow cytometry (**Figure S1B)**, indicative of their endothelial identity. We measured the proliferative index of hiPSC-derived ECs by IF staining for Ki67 and DAPI. rBMECs proliferated at a similar rate to bpECs, but at a lower rate than primary HBMECs when cultured without growth factors (VEGF-A and bFGF; **Figure 1I)**. To assess the response to angiogenic stimuli, we performed a wound-scratch assay on EC monolayers (bpECs, rBMECs and HBMECs) treated with either basal media or media supplemented with an angiogenic cocktail [VEGF-A, bFGF, and hepatocyte growth factor (HGF], analyzed the area of wound closure during a 36-hour time lapse as a readout of cell migration and calculated the angiogenic potential (see Methods for details; **Figure 1I-K; S1K-L**). By 36 hours, rBMEC migration increased in response to the angiogenic cocktail compared to control conditions (**Figure 1J**). Primary HBMECs and rBMECs exhibited a comparable angiogenic potential, while bpECs responded more strongly to the angiogenic stimuli (**Figure 1K**). Overall, these data demonstrate that rBMECs express both key EC and BBB markers and proliferate and migrate in response to angiogenic stimuli.

### Transcriptome profiling reveals that rBMECs express a subset of the BBB gene repertoire

We used bulk RNA sequencing (bulk RNA-seq) to characterize the CNS EC identity and BBB gene expression in rBMECs. We used principal component analysis (PCA) to compare the rBMEC transcriptome to other human cell types from published datasets^28,37–39^. rBMECs clustered most closely with ECs and furthest apart from choroid plexus cells, epithelial cells, and iBMECs^28^ (**Figure 2A; Table S1A**). Together with our IF staining and flow cytometry data, this PCA plot confirms that rBMECs have an endothelial identity. The PCA comparison of rBMECs with other EC populations showed that they cluster closer to HBMECs and further away from non-CNS ECs **(Figure 2B; Table S1B).** We performed differential gene expression between rBMECS and HBMECs, or bpECs, to identify which CNS EC and BBB gene signatures are distinct between these populations. A significant number of CNS ECs and BBB genes were distinct between rBMECs and HBMECs (**Figure 2C; Table S1C-D**) or bpECs (**Figure S2B; Table S2C, E**), although this difference was higher with HBMECs. Gene ontology (GO) analysis of the differentially expressed genes (DEGs) revealed that pathways related to “blood vessel and vasculature development”, “regulation of EC migration” and “BBB permeability maintenance” were upregulated in rBMECs compared to HBMECs **(Figure 2D)**. Several BBB genes encoding for TJs (e.g. CLDN5, OCLN, JAM2), transporters (e.g. SLC2A1, ABCC4, and SLC1A1), receptor-mediated transcytosis (e.g. CAV1, CAV2, LRP10) and ECM components (e.g. COL4A1, COL4A2) were higher in rBMECs compared to HBMECs (**Figure 2E-G**; red); however other BBB genes within these categories were either not significantly different or lower in expression (**Figure 2E-G**; blue). In addition, rBMECs expressed higher levels of CNS EC identity markers (e.g. FOXF2, FOXQ1, SPOCK2, ITM2A, PROM1, LRP8, and APCDD1) compared to primary HBMECs (**Figure 2H; S2A**). Although rBMECs shared a similar gene expression profile with bpECs (**Figure S2B**), several key BBB genes (e.g. ITIM2A, SPOCK2, CLDN5, OCLN, TJP2, SLC2A1 and COL4A1) were upregulated significantly in rBMECs compared to bpECs **(Figure S2C-G)**, suggesting improvement in BBB gene signatures. Moreover, rBMECs showed reduced expression of LAMs (e.g. VCAM1 and SELP) compared to either HBMECs (**Figure 2G**), or bpECs (**Figure S2E**), indicative of a CNS EC identity. In summary, the rBMEC transcriptome analysis shows stronger expression of several CNS EC identity and BBB gene signatures compared to primary HBMECs and bpECs.

**Figure 2.**
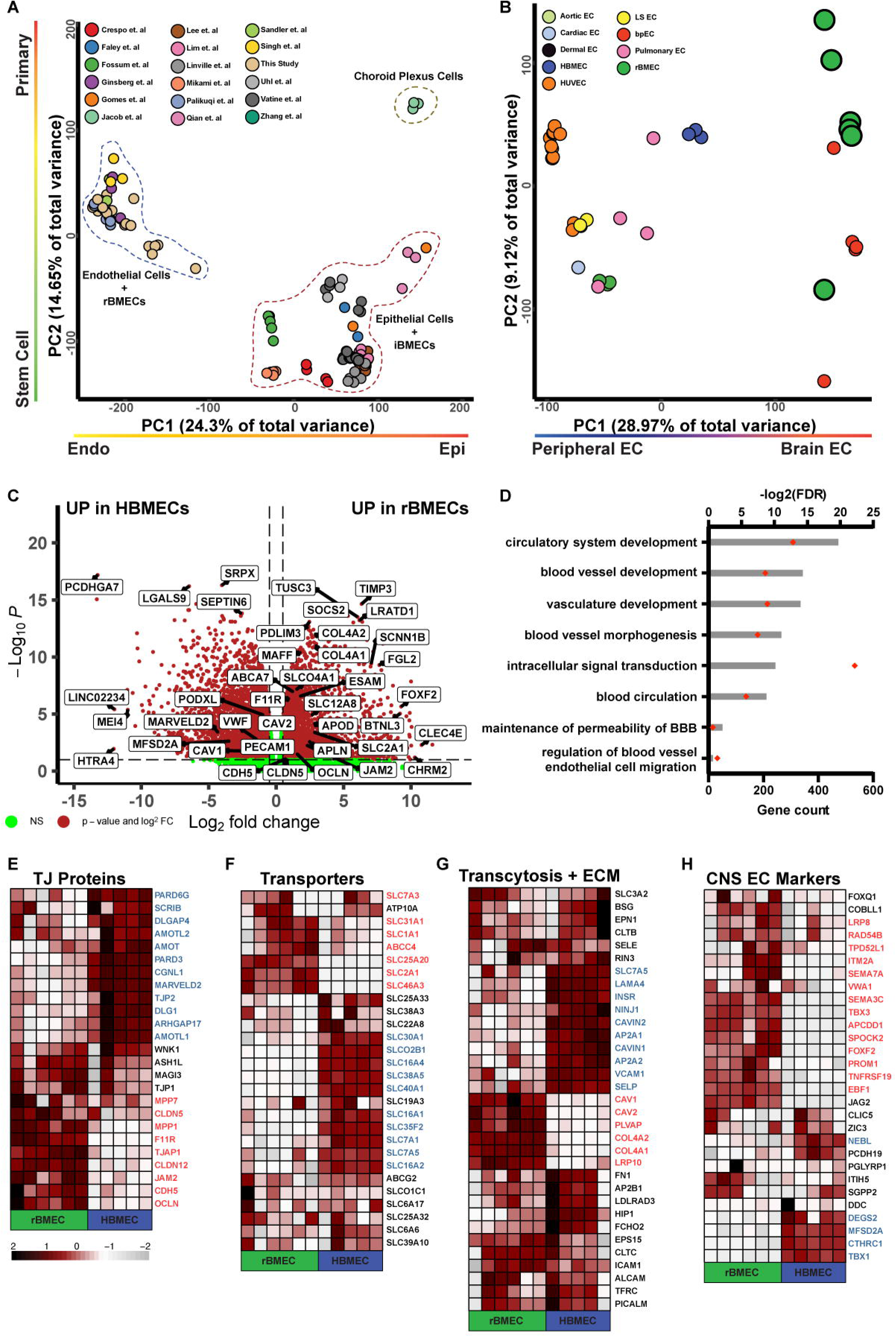
The rBMEC transcriptome shows higher expression of BBB molecular identity genes compared to primary HBMECs. **A)** PCA plot illustrating endothelial vs. epithelial identity of 155 sequenced RNA samples from several published studies and this study. **B)** PCA plot illustrating peripheral vs. brain EC identity of 58 differentiated and primary EC samples from published studies and this study. **C)** Volcano plot highlighting significantly up- and downregulated BBB-specific genes in rBMECs compared to HBMECs. **D**) Upregulated GO analysis pathways in rBMECs compared to HBMECs. **E-H)** Heatmap of BBB-specific genes (TJ proteins, Transporters, Bulk and Receptor-Mediated Transcytosis, Extracellular Matrix, and CNS EC genes) between rBMEC and HBMECs. The upregulated genes in rBMECs are shown in red and the upregulated genes in HBMECs are shown in blue.

### rBMECs have robust BBB functional properties

We next investigated the functional properties of rBMECs in comparison to bpECs and primary HBMECs. First, we measured TEER over time as a readout of the paracellular EC barrier formed by tight junctions. rBMECs differentiated from the IMR90 line showed a higher average TEER (40.5 ± 4.2 Ω*cm^2^) compared to either bpECs (25.0 ± 0.4 Ω*cm^2^), or primary HBMECs (24.7 ± 3.0 Ω*cm^2^), for >300 hours. The area under the curve (AUC) for the last 100 hours of recording was two-fold higher in rBMECs compared to HBMECs (**Figure 3A, B**). TEER was also increased in rBMECs derived from the other hiPSC lines FA10, NCRM5, and NCRM4 compared to primary HBMECs (**Figure S3A, B, E, F, H, I**). Next, we used three different molecular weight tracers, sodium fluorescein (NaF = 376 Da), 3- and 70-kDa dextran to assess their diffusion (permeability co-efficient) across the EC monolayer in a Transwell system. rBMECs showed a significantly reduced permeability co-efficient (Pe) for the smallest tracer (NaF) across the monolayer compared to bpECs and HBMECs indicative of tighter paracellular barrier **(Figure 3C)**. Although HBMECs showed a significantly higher Pe compared to rBMECs for the medium size tracer (3kDa Dextran), there was no difference between rBMECs and bpECs **(Figure 3D)**. In contrast, the Pe for the 70 kDa Dextran was very low across all three EC lines (rBMECs, bpECs and hBMECs) suggesting very low transcellular transport **(Figure S3C, D).** Overall, the TEER and tracer permeability data demonstrate that rBMECs have a tighter paracellular barrier compared to primary HBMECs and bpECs.

**Figure 3.**
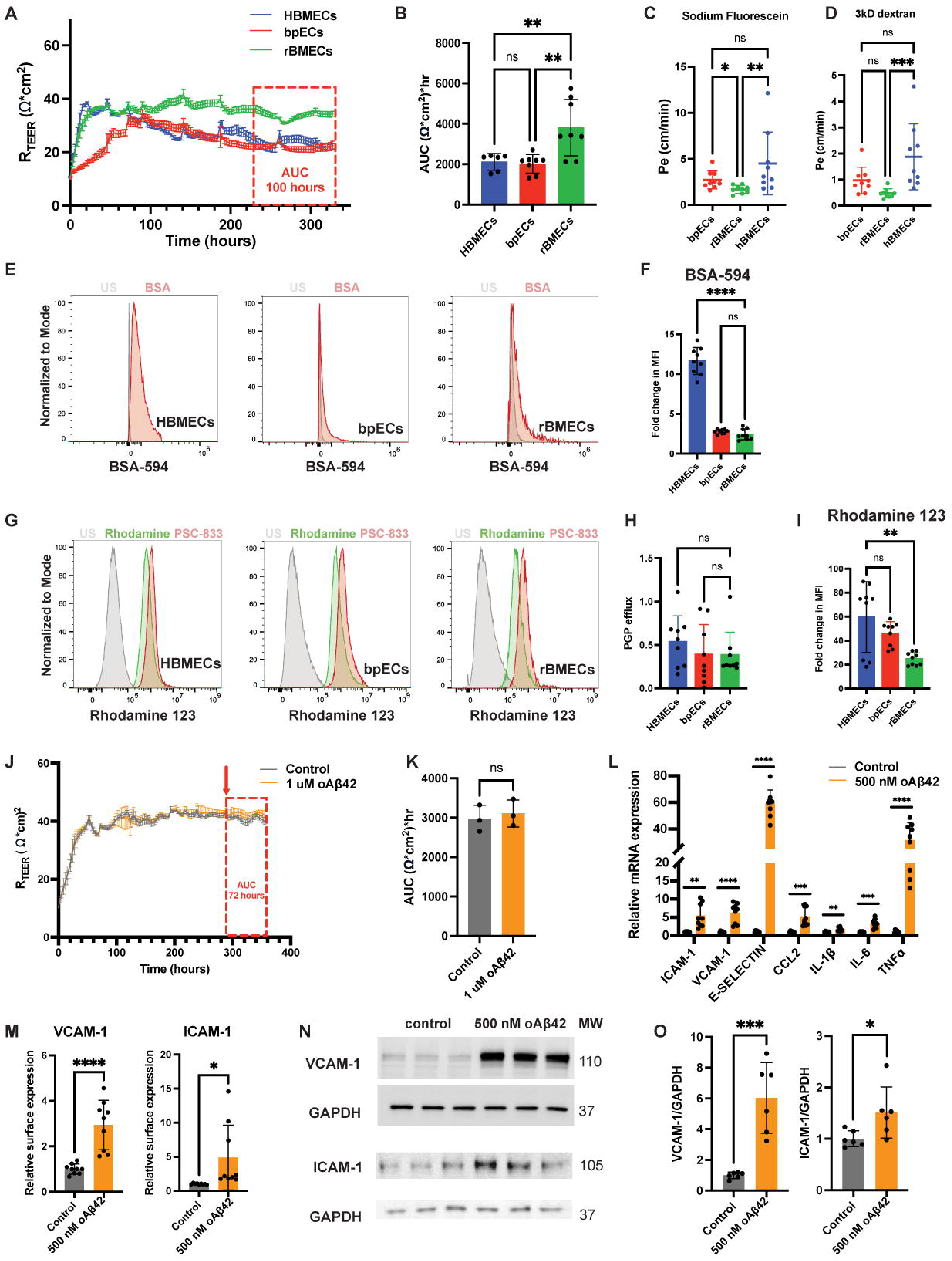
rBMECs have strong functional BBB properties and respond to inflammatory stimuli. **A-B)** Representative ECIS TEER recording and dotted bar graph of the area under the curve (AUC) quantification for IMR90 hiPSC-derived bpECs, rBMECs, and primary HBMECs. The red box indicates the period used for the AUC quantification. Mean ± SD, n = 6-8 from 3-4 experiments, one-way ANOVA, **p<0.01. **C-D)** Dotted bar graphs of the transwell permeability across EC monolayers to sodium fluorescein (NaF) and 3kDa dextran. Mean ± SD, n = 9 from three experiments, ANOVA with Kruskal-Wallis Test, *p<0.05, **p<0.01, ***p<0.001. **E-F)** Representative histograms and quantification of BSA-AF594 uptake in HBMECs, bpECs and rBMECs. Mean ± SD, n = 9 from three experiments, one-way ANOVA, ****p<0.0001. **G-I)** Representative histograms of Rhodamine123 MFI and PGP efflux in HBMECs, bpECs and rBMECs. n = 9 from 3 independent experiments, one-way ANOVA, **p<0.01. **J-K)** Representative ECIS TEER and dotted bar graph of the AUC quantification for rBMECs during 72 hr treatment with F12 vehicle or 1 uM oAβ42 (red box), n = 9-10 from three experiments, unpaired t-test, ns p>0.05. **L)** RT-qPCR for several inflammatory genes in rBMECs after 6 hours of 500nM oAβ42 treatment; n = 8-9 from three experiments, unpaired t-test, **p<0.01, ***p<0.001, ****p<0.0001. **M)** VCAM-1 and ICAM-1 surface expression quantification after 6 hours 500nM oAβ42; n = 6 from two experiments, unpaired t-test, **<0.01, ****<0.0001. **N-O)** Representative western blots and quantification for VCAM-1 and ICAM-1 after 6 hours of 500nM oAβ42; n = 6 from two experiments, unpaired t-test, ***<0.001.

To assess transcellular rBMEC barrier properties, we examined caveolar-mediated transcytosis by incubating EC monolayers with a fluorescently-tagged bovine serum albumin (BSA; MW = 60 kDa) and measuring its uptake inside cells by flow cytometry. rBMECs and bpECs showed very small changes in the mean fluorescence intensity (MFI) of the intracellular fluorescent BSA compared to HBMECs (**Figure 3F**), indicative of low caveolae-mediated transcytosis. Finally, we assessed the function of P-GLYCOPROTEIN (PGP, ABCB1), a transporter that effluxes drugs from the brain to the blood^40^. We incubated HBMECs, bpECs, and rBMECs monolayer cultures with Rhodamine 123, which is effluxed by PGP^41^, in the presence or absence of a PGP-specific inhibitor, PSC-833. Cells were subsequently analyzed by flow cytometry to assess changes in MFI of the dye. We calculated the efflux rate based on the MFI change in the presence / absence of the inhibitor. HBMECs, bpECs, and rBMECs had comparable efflux rates in response to PSC-833, suggesting similar PGP activity **(Figure 3G, H**). However, rBMECs had a lower fold change in Rhodamine 123 MFI compared to HBMECs (**Figure 3I**) suggesting an overall greater efflux. These data suggest that rBMECs have an increased efflux of rhodamine compared to bpECs and HBMECs, although PGP activity is similar.

We investigated the ability of rBMECs to respond to pro-inflammatory and disease-relevant stimuli such as oligomeric amyloid-β (1-42) peptides (oAβ42). Most AD patients have amyloid deposits in the cerebral vasculature in post-mortem brains studies^42–44^, decreased microvascular density, vBM thickening, BMEC and pericyte damage^45^, inflammatory changes in vessels, and microglial activation around blood vessels^46^. We first assessed if oAβ42 affected the paracellular barrier properties of rBMECs. Contrary to published studies^47^, there was no change in TEER over 72 hours when rBMECs were exposed to either 500 nM or 1 uM oAβ42 (**Figure 3J, K**; data not shown). In contrast, the pro-inflammatory cytokines TNFα and IL1β significantly reduced the rBMEC TEER over 24 hours **(Figure S3J, K),** confirming that rBMECs respond to pro-inflammatory cytokines. To determine if oAβ42 induces EC activation, we extracted mRNA and measured changes in LAMs and chemokines via qPCR. *VCAM-1* was increased by 5.2-fold, and *ICAM-1* was increased by 1.7-fold after 24 hours of 1uM oAβ42 exposure (**Figure S3G**). Chemokine ligand 2 (*CCL2*), a major cytokine implicated in the pathogenesis of ischemic stroke^48^ and other BBB-compromising pathologies^49,50^ and a powerful monocyte and T-cell attractant^48^, was also upregulated in rBMECs treated for 24 hours with 1 uM oAβ42 (**Figure S3G**). Therefore, rBMECs respond to 1 uM oAβ42 by upregulating expression of LAMs and pro-inflammatory cytokines. To examine acute effects of oAβ42, we treated rBMECs with 500 nM oAβ42 for 6 hours and analyzed changes in LAMs and pro-inflammatory cytokines by qPCR, western blot, and flow cytometry, as possible downstream effectors of oAβ42-mediated dysfunction. oAβ42 significantly increased mRNA expression of *ICAM-1* and *VCAM-1, E-SELECTIN, CCL2, IL-1β*, *IL-6*, and *TNFα* within 6 hours (**Figure 3L**). Additionally, ICAM-1 and VCAM-1 total and surface protein levels as measured by western blot and flow cytometry, respectively, were upregulated in rBMECs after acute treatment with 500 nM oAβ42 (**Figure 3M-O**). These data are consistent with recent findings that primary HBMECs upregulate *VCAM-1, ICAM-1, E-SELECTIN*, and *CCL2* after treatment with conditioned media from hiPSC-derived cortical neurons harboring the APP^Swe/Swe^ mutation (KM670/671NL)^51^. Finally, we sought to identify potential signaling pathways that might contribute to oAβ42-induced inflammatory phenotypes in rBMECs. Phosphorylation of signal transducer and activator of transcription 3 (STAT3), a key mediator of proinflammatory processes in brain ECs^52,53^, was elevated in rBMECs treated with oAβ42 **(Figure S3L-M).** Therefore, β-amyloid peptides act as pro-inflammatory stimuli in rBMECs to upregulate a subset of EC inflammatory gene repertoire that may contribute to increased interactions with immune cells. Overall, these functional studies demonstrate that rBMECs have strong functional BBB properties and are suitable to study immune - EC interactions and the potential impact of disease-relevant stimuli.

### Modeling the human 3D NVU/BBB using rBMECs and hiPSC-derived astrocytes and pericytes

We next investigated whether rBMECs can interact with iPSC-derived astrocytes and pericytes and generate a fully iPSC-derived human neurovascular unit (NVU) in a 3D microfluidic system (**Figure 4A**). We generated iPSCs-derived neural crest pericytes (iPCs) following an established protocol, where iPSCs were treated with a neural crest differentiation media supplemented with CHIR99021^54^. iPCs were later sorted on CD146^+^/PDGFRβ^+^ to ensure a homogeneous population prior to introduction into the 3D NVU microfluidics system (**Figure S4A**). iPCs expressed PDGFRβ and NG2 (**Figure S4C**) and they had a similar transcriptome profile with primary human pericytes and iPSC-derived neural crest pericytes from prior studies^55^ (**Figure S4E**). hiPSCs-derived astrocytes (iACs) were generated from iPSC-derived neural progenitor cells (NPCs) as described^56^ with the addition of NFIA transduction of NPCs. iACs expressed high levels of astrocytic proteins GFAP and VIMENTIN (**Figure S4D**) and they were similar at the transcriptome level with primary human astrocytes (**Figure S4F**).

**Figure 4:**
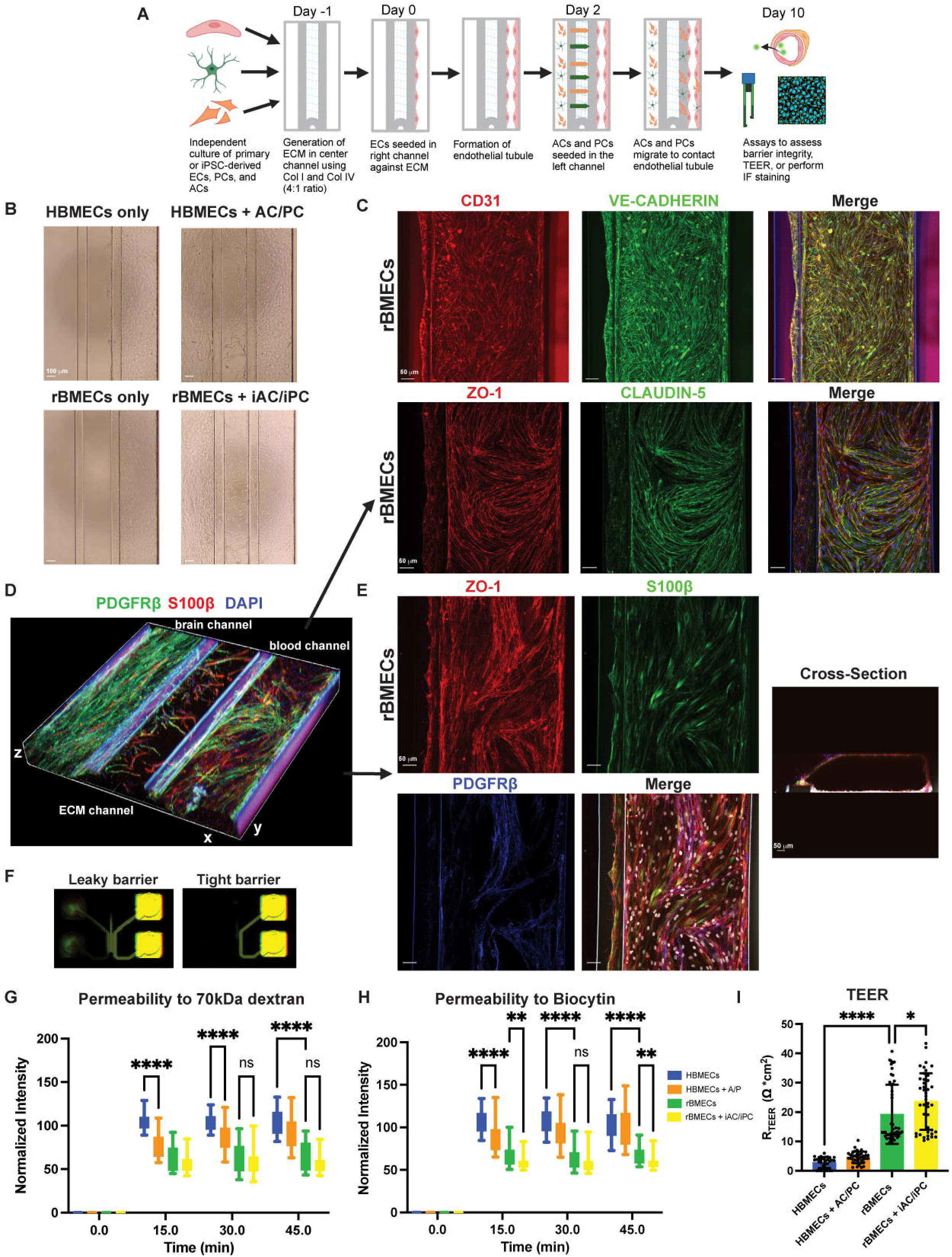
rBMECs tubules have improved barrier properties when co-cultured with iPSC-derived astrocytes and pericytes in a 3D microfluidic NVU system. **A)** Schematic diagram for generating the 3D NVU/BBB microfluidic system. **B)** Brightfield images of EC tubules formed by HBMECs and rBMECs with (right) and without (left) primary or iPSC-derived astrocytes and pericytes, respectively. **C)** Representative IF staining of rBMEC tubules for CD31 and VE-CADHERIN (top) or ZO-1 and CLAUDIN-5 (bottom). Scale bar = 50 µm **D)** 3D image of IF staining within the 3D NVU microfluidics system showing iPCs (PDGFRβ) and iACs (S100β) migrating from the “brain” to the “blood” channel where the rBMECs tubules are located. **E)** Representative triculture staining with rBMECs (ZO-1), iACs (S100β), and iPCs (PDGFRβ) in the blood channel; cross sectional view shows tubule with a lumen. **F)** Representative image showing diffusion of fluorescent tracers through leaky and tight tubules. **G-H)** Box plots showing the normalized intensity of 70 kDa dextran and biocytin (890 Da) tracers in the middle channel indicative of permeability; n = 36-41 NVU chips from three experiments. two-way ANOVA, **p<0.01, ****p<0.0001. **I)** Dotted bar graph of the TEER measurement in the 3D NVU/BBB microphysiological system using the OrganoTEER for EC tubules alone or together with PCs/ACs; n = 30-48 chips over three experiments, one-way ANOVA, ****p<0.0001, *p<0.05.

For the 3D microfluidic system, we used the MIMETAS 64-chip OrganoPlate® comprised of three lanes^57,58^. rBMECs or HBMECs were introduced on Day 0 in the right “blood channel” against a collagen I/IV gel mixture loaded in the middle “ECM” channel, to enable tubule formation. iACs and iPCs were introduced in the left “brain” channel after 2 days and the system was allowed to mature for 10 days with bidirectional flow (**Figure 4A**). By 10 days, rBMECs formed continuous tubules and iACs and iPCs migrated towards the “blood vessel” channel to contact the EC tubules (**Figure 4B**). rBMEC tubules were positive for the EC markers CD31 and VE-CADHERIN and TJ proteins CLAUDIN-5 and ZO-1 (**Figure 4C; Movies S1, S2**). In 3D co-cultures of rBMECs with iACs and iPCs, ZO-1^+^ ECs were in close proximity to S100β^+^ (astrocyte) and PDGFRβ^+^ (pericyte) cells, suggesting that iACs and iPCs migrated across the ECM channel to contact the rBMEC tubule (**Figure 4D, E; Movies S3, S4**). Therefore, rBMECs can form tubules and interact with iPSC-derived astrocytes and pericytes to generate a 3D human NVU.

To assess BBB properties of rBMECs in the 3D microfluidic system, we performed tracer diffusion studies using both small (biocytin, 890Da) and large (dextran, 70kDa) tracers. We introduced the tracers into the blood channel, allowed them to diffuse across the chip for 45 minutes, and measured the mean fluorescent intensity in the middle channel as a readout of tracer leakage through the tubule at three distinct time points (15, 30 and 45 minutes). rBMECs exhibited a significant lower leakage of biocytin and 70kDa dextran compared to primary HBMECs, suggesting that they had tighter paracellular and transcellular barrier properties (**Figure 4F-H**). Moreover, rBMECs cocultured with iACs and iPCs exhibited lower permeability to biocytin compared to rBMECs alone by 45 minutes, suggesting that iPCs/iACs increase the barrier properties of rBMECs (**Figure 4H**). Finally, we measured TEER across the EC tubule in the 3D microfluidics system using the OrganoTEER instrument from MIMETAS^57^. While there was considerable variability across experiments, rBMEC tubules displayed on average a six-fold higher TEER compared to HBMEC tubules alone (**Figure 4I, S4G**). rBMEC tubules also had a significantly higher TEER when co-cultured with iACs/iPCs (**Figure 4I, S4G**) consistent with the tracer diffusion studies (**Figure 4H**). Thus, rBMECs form stable and continuous tubules in the MIMETAS 3D microfluidic system, and they interact well with iPCs/iACs to form an iPSC-based NVU with robust BBB properties. This 3D system resembles closely the *in vivo* BBB at both structural and functional levels and can be used to study either NVU - immune dysregulation or the crosstalk between genetic and environmental factors critical for pathogenesis of many neurological diseases.

## DISCUSSION

There generation of an *in vitro* human NVU system with robust BBB properties to model neurological diseases and develop effective therapeutics to enter the CNS has been a major technological hurdle for many decades^14–17^. While several human iPSC-derived BBB models have been developed in the last decade, their utility has been limited by a lack of brain-specific organotypic EC identity and an inability to replicate the multifaceted properties of the BBB. Current methods have either generated ECs with limited BBB properties^30,35^, weak immune responses^22,23,26,27,30^, generic but not brain-specific cell identity^28,59^, or mixed epithelial-endothelial identity^21–24,26,27,60^. In this study, we show that we can generate ECs with robust brain-specific identity and BBB properties (rBMECs) through a combinatorial strategy to initially differentiate hiPSCs into bpECs followed by overexpression of two brain-specific TFs FOXF2 and ZIC3 to drive brain organotypicity. rBMECs express higher levels of several key BBB markers belonging to tight junctions, transcytosis, transporters and key CNS EC identity markers at both transcript and protein levels, are capable of undergoing angiogenesis, and respond robustly to oAβ42 treatment (i.e. inflammatory response). In addition, rBMECs have a tighter paracellular barrier, reduced caveolae-mediated transcytosis, and equivalent PGP activity compared to primary HBMECs at a functional level. rBMECs are able to make contacts with iPSC-derived pericytes and astrocytes in a 3D system with flow and strengthen their barrier properties through mutual cell-cell interactions with these other NVU cells, mirroring their 3D organization in the human brain. While several static^61,62^ and perfusable^21,24,63,64^ iPSC-derived 3D NVU models have been published, our 3D model uses brain-specific ECs that have the capacity to model multiple properties of the human BBB. Overall, our data demonstrate that rBMECs generated through this new approach can be utilized as an *in vitro* human-specific BBB monolayer model and/or a 3D microfluidic NVU model in combination with iPSC-derived PCs/ACs.

In contrast to our approach of overexpressing FOXF2 and ZIC3 simultaneously, Roudnicky et al. have overexpressed either FOXF2 or ZIC3 alone in ECs generated via a different protocol^59^ through adenoviral transduction. FOXF2 overexpression seems to improve some of the BBB properties of ECs by increasing *OCLN* mRNA expression by qRT-PCR, and upregulating Wnt-β-catenin signaling through pathway analysis^65^. FOXF2 also upregulated the inflammatory response pathway. In contrast, ZIC3 overexpression had no effect on the barrier properties of ECs^65^. We find that FOXF2 / ZIC3 simultaneous overexpression is sufficient to increase rBMEC barrier properties, but does not induce inflammatory LAMs; *SELP* and *VCAM1* were downregulated, whereas *SELE* and *ICAM1* were unaltered in rBMECs compared to primary HBMECs. Our findings suggest that a FOXF2 / ZIC3 combination is critical for the acquisition of a larger repertoire of BBB properties including barrier strength, suppression of transcytosis genes and suppression of the inflammatory LAMs. However, the precise mechanism by which FOXF2/ZIC3 reprogram bpECs to strengthen BBB characteristics remains unclear. We postulate that bpECs generated via the Praca et al. protocol^35^ are more amenable to acquire stronger BBB properties than other EC cell types, as our preliminary efforts to transduce either endothelial progenitor cells^66^ or ECs generated from a commercial kit (StemCell Technologies) with FOXF2/ZIC3 failed to generate cells with similar BBB properties as rBMECs. Therefore, FOXF2 and ZIC3 reprogramming seems most effective in a more “plastic” environment present in the BBB-primed EC which already has some BBB-like properties.

How does Wnt/β-catenin signaling activation affect the BBB repertoire in rBMECs? We have found that administration of CHIR99021, a potent activator the Wnt/β-catenin signaling does not have an additive effect on BBB properties or “brain identity” of rBMECs (data not shown). This was somewhat surprising since a recent study showed that a small molecule cocktail (cARLA) that activates Wnt-β-catenin and cAMP signaling and inhibites TGF-β signaling in generic ECs^67^ was sufficient to induce robust BBB properties (tight barrier, high efflux activity) and expression of a subset of brain EC genes^67^. However, cARLA failed to increase expression of *FOXF2*, *ZIC3*, and some key BBB genes (e.g. *ITM2A*, *OCLN*) which are robustly upregulated in our rBMECs. cARLA also increased expression of the efflux transporter *ABCB1*, which was not increased by FOXF2 and ZIC3 overexpression, despite comparable levels of PGP efflux activity with primary HBMECs. These two diverse strategies (cARLA vs. FOXF2/ZIC3 reprogramming) can induce complementary BBB properties, whereas FOXF2 and ZIC3 reprogramming is critical to upregulate essential “brain identity” markers that cannot be achieved by the small molecule cocktail method. Future studies will determine whether a combination of cARLA and FOXF2/ZIC3 reprogramming strategies can achieve both superior BBB properties and brain organotypicity in ECs.

Ultimately, one of our goals is to generate an *in vitro* human NVU/BBB model that could faithfully model pathogenic mechanisms of neurological diseases. We investigated the ability of rBMECs to respond to pro-inflammatory and disease-relevant stimuli such as oligomeric amyloid-β (1-42) peptides (oAβ42) elevated in AD patients. In contrast to published studies^68^, oAβ42 did not affect barrier tightness of rBMECs, but was sufficient to upregulate significantly expression of pro-inflammatory cytokines and LAMs, including *VCAM-1* and *ICAM-1*. Here, we used a lower, more physiologically relevant dosage of oAβ42 than previously described^68^, which is non-toxic in our hands and may explain the lack of effect on the barrier properties. This also suggests that oAβ42, unlike fibrillar forms of Aβ, does not simply damage the BBB to cause dysfunction, but might rather serve as an immune signaling molecule, consistent with the antimicrobial protection hypothesis of AD^69^. Taken altogether, we have established a molecularly and functionally faithful brain EC cell type that can be widely either alone or with other NVU cells to study pathogenic mechanisms of neurological diseases and drug delivery to the CNS.

### Limitations

While rBMECs have BBB molecular identity and functional properties, they do not fully recapitulate all aspects of the human BBB. For instance, they do not robustly express the full repertoire of BBB-specific transporters (e.g. MSFD2A). In addition, one major drawback of the differentiation strategy is the low efficiency to produce bpECs (Sort A). Unfortunately, our attempts to reprogram ECs generated from other protocols failed to produce cells with robust BBB properties. Finally, we have not explored the mechanisms by which FOXF2 and ZIC3 drive and sustain BBB properties in rBMECs. This will be a target of future studies.

## ACKNOWLEDGEMENTS

We thank Raphael Lis for providing FOXF2 and ZIC3 expression plasmids, and Paroma Malick in Alex Chavez’s lab for subcloning FOXF2 and ZIC3 into pLEX_307. Flow cytometry and cell sorting experiments were performed in the Columbia Stem Cell Initiative Flow Cytometry core facility under the leadership of Michael Kissner. This work was supported by NIH grants from the Heart Lung and Blood Institute [R61/R33 HL159949 and R61/R33 HL159949 (PB)] and National Institute of Aging [RF1AG078352 and RF1AG078352 (PB)]. UA, and DA are also supported by grants from the National Eye Institute (R01EY033994), National Institute of Neurological Disorders and Stroke (R21NS130265), the International OCD Foundation and the Global Lyme Alliance. AAS is also supported by the Henry and Marilyn Taub Foundation and Thompson Family Foundation (TAME-AD).

## AUTHOR CONTRIBUTIONS

AC, RP, and PB performed experiments, analyzed data, and drafted the manuscript. UA performed the bioinformatic analysis. ER performed some experiments and provided invaluable technical support. PBD performed experiments and analyzed data. AAS and DA conceptualized the project, provided the funding, oversaw all work, and drafted the manuscript. All authors edited the manuscript.

## STAR* METHODS

### KEY RESOURCE TABLE

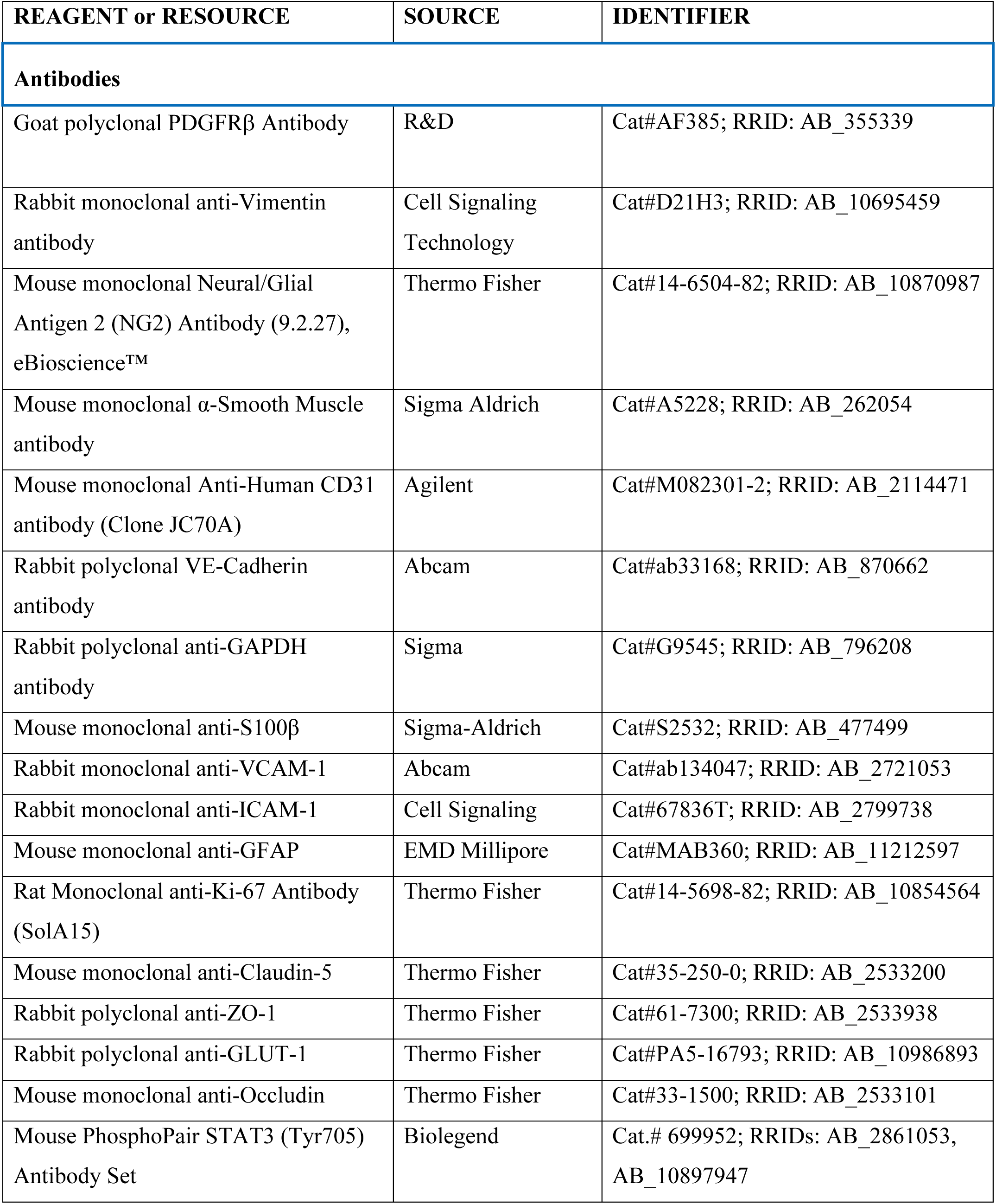
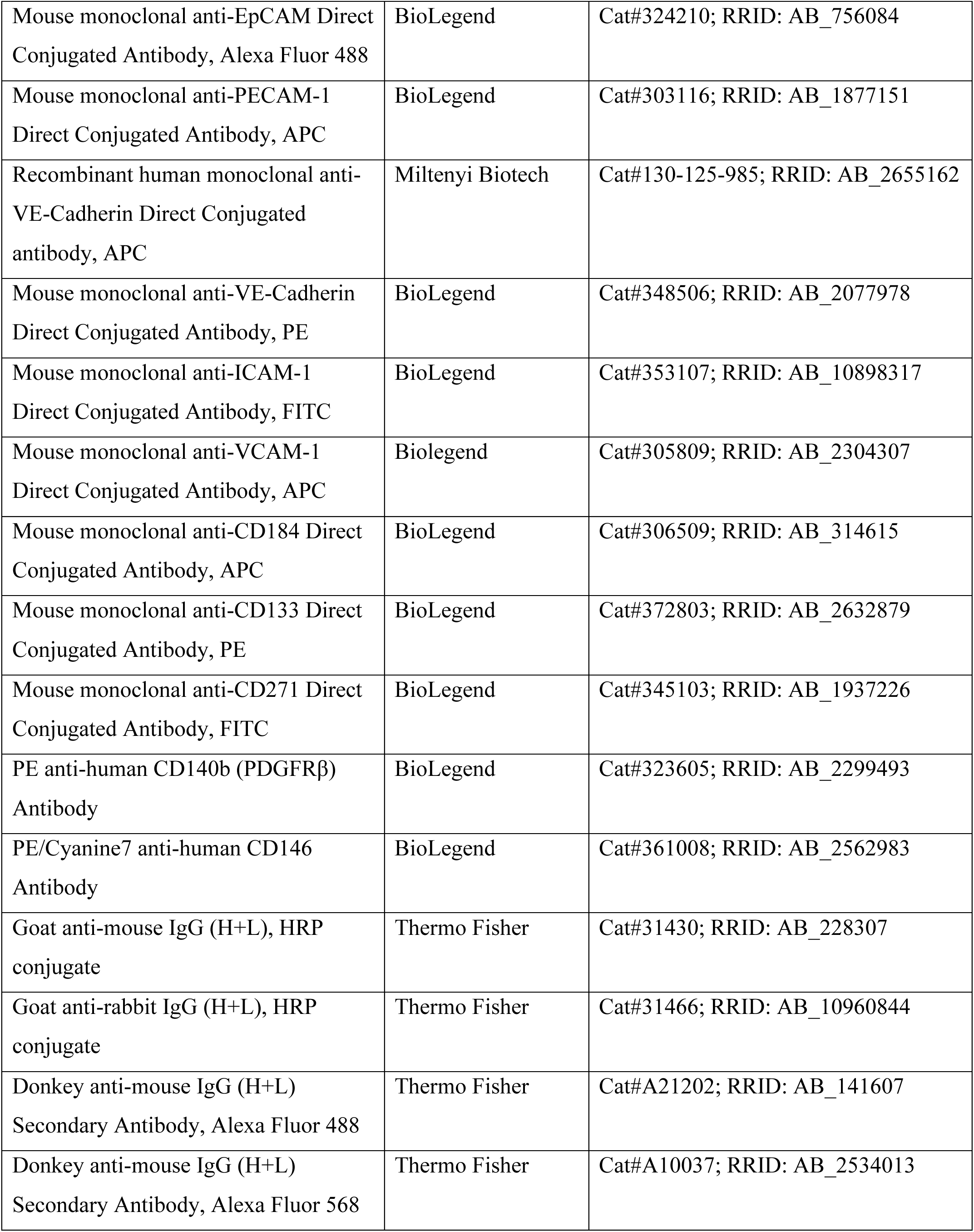
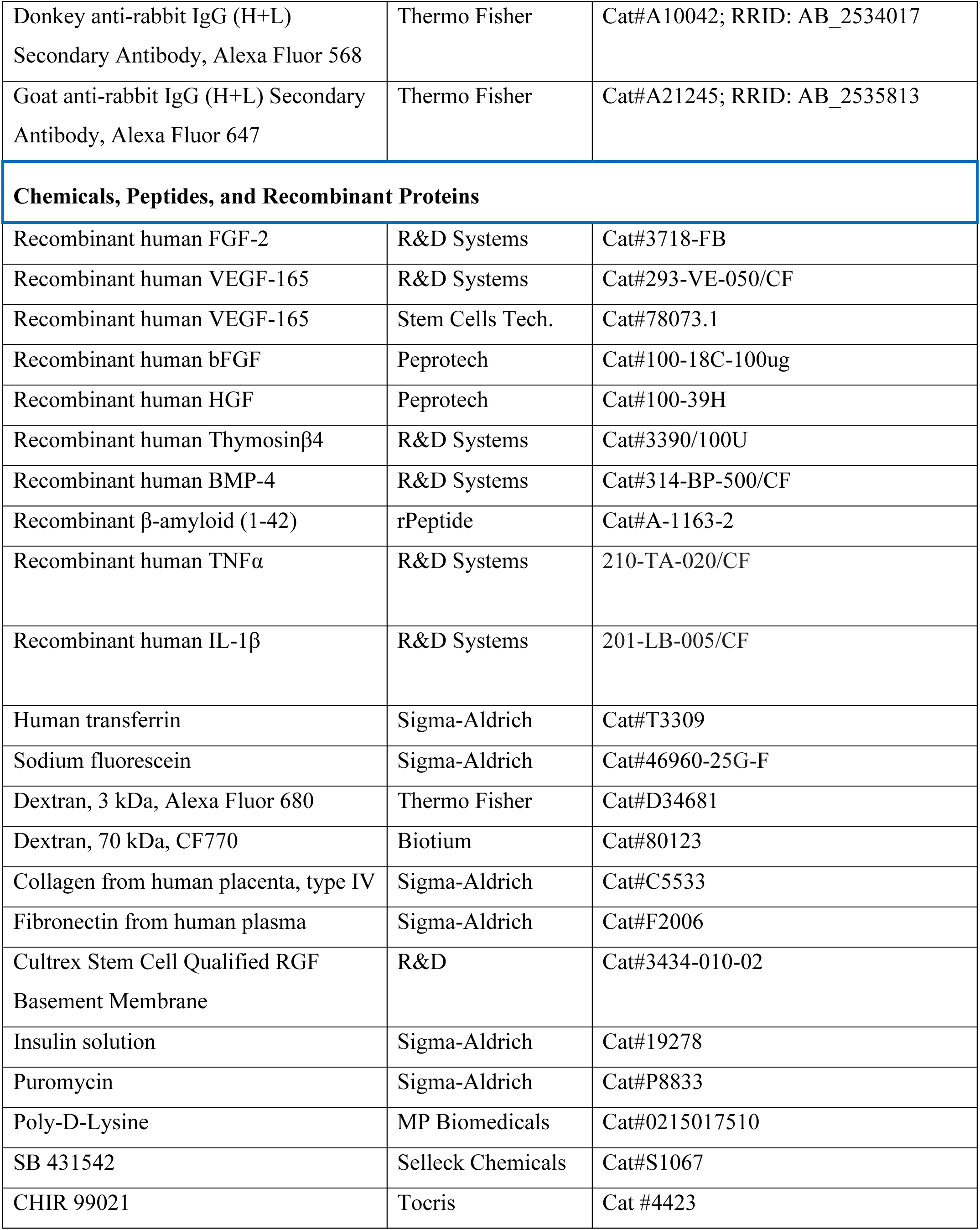
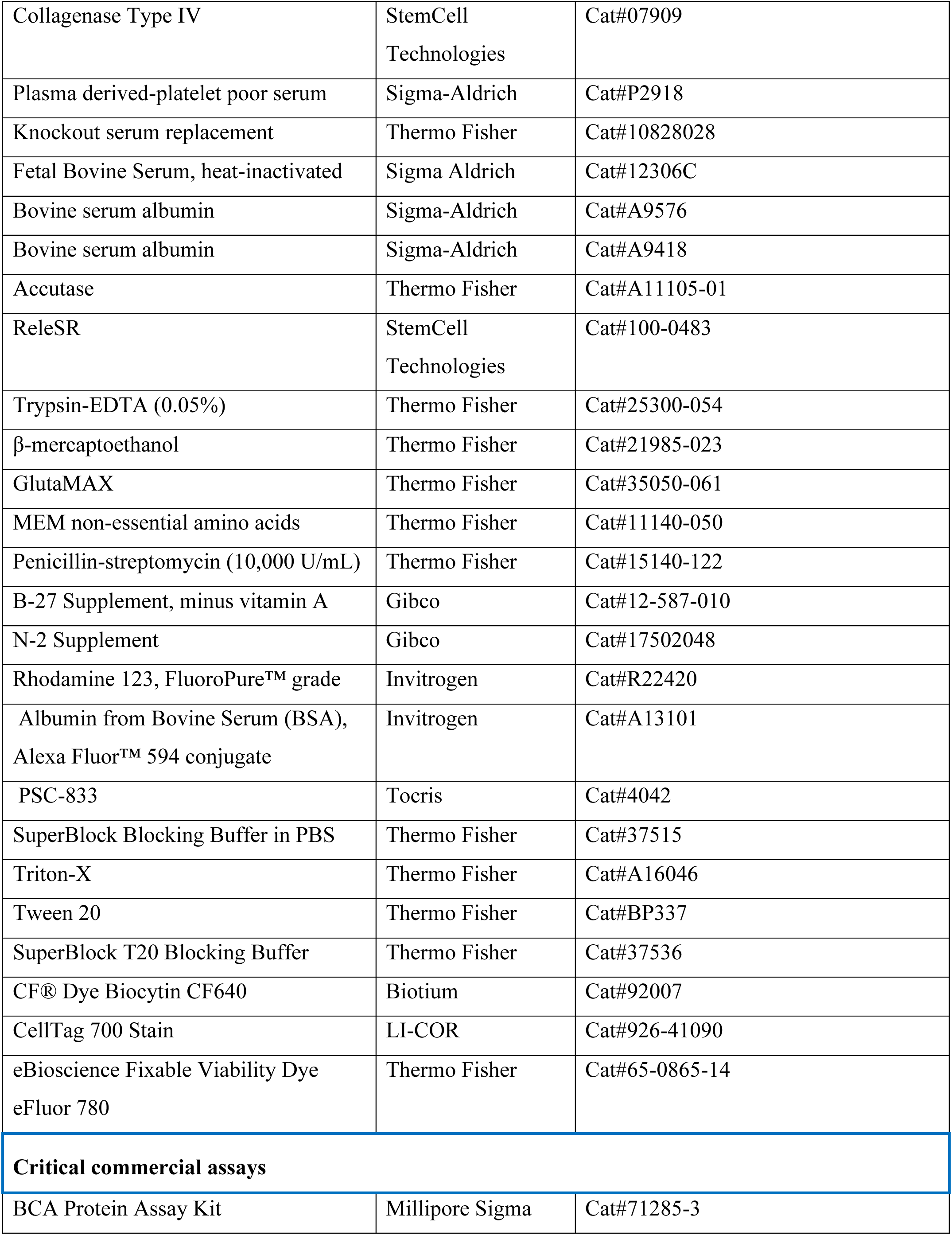
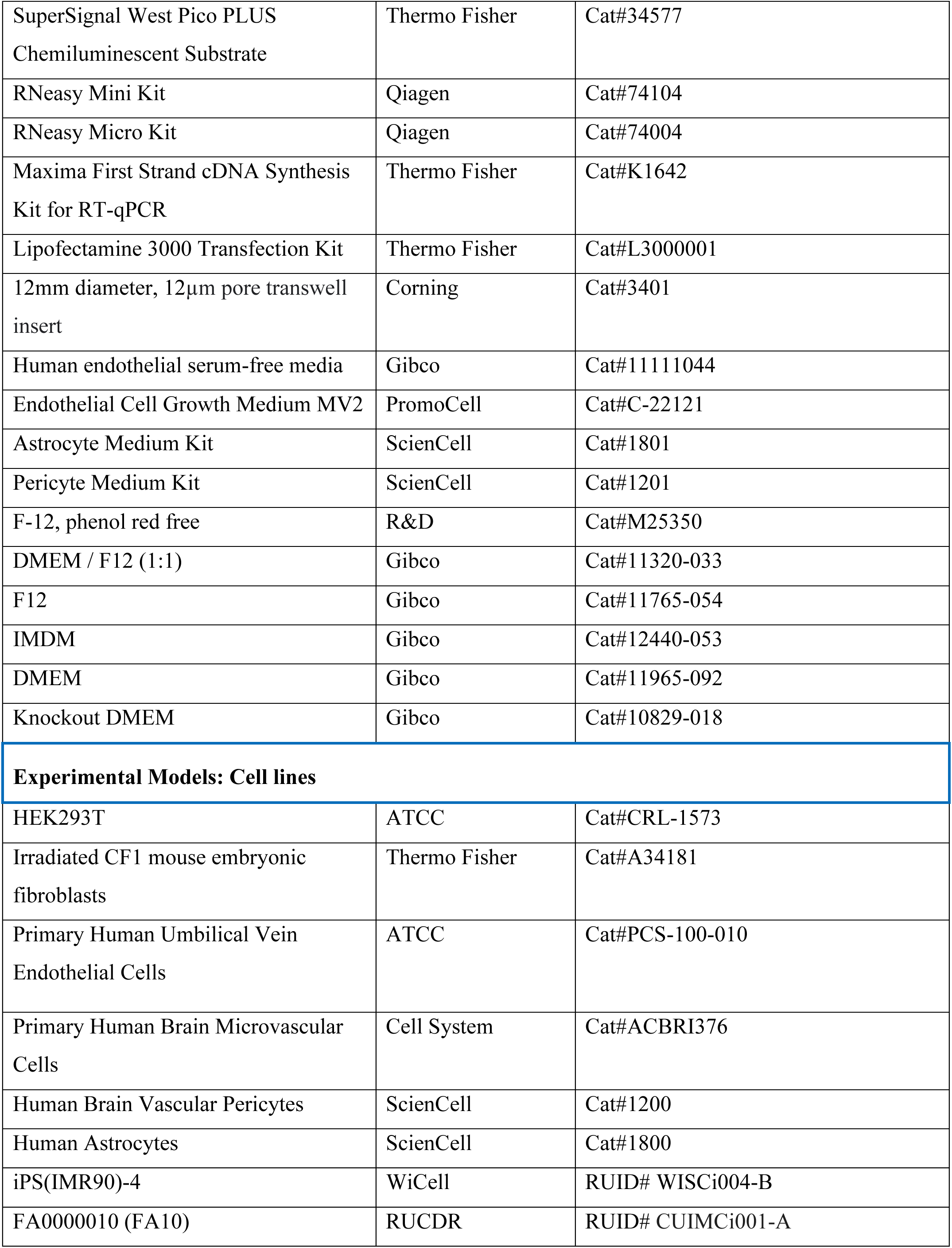
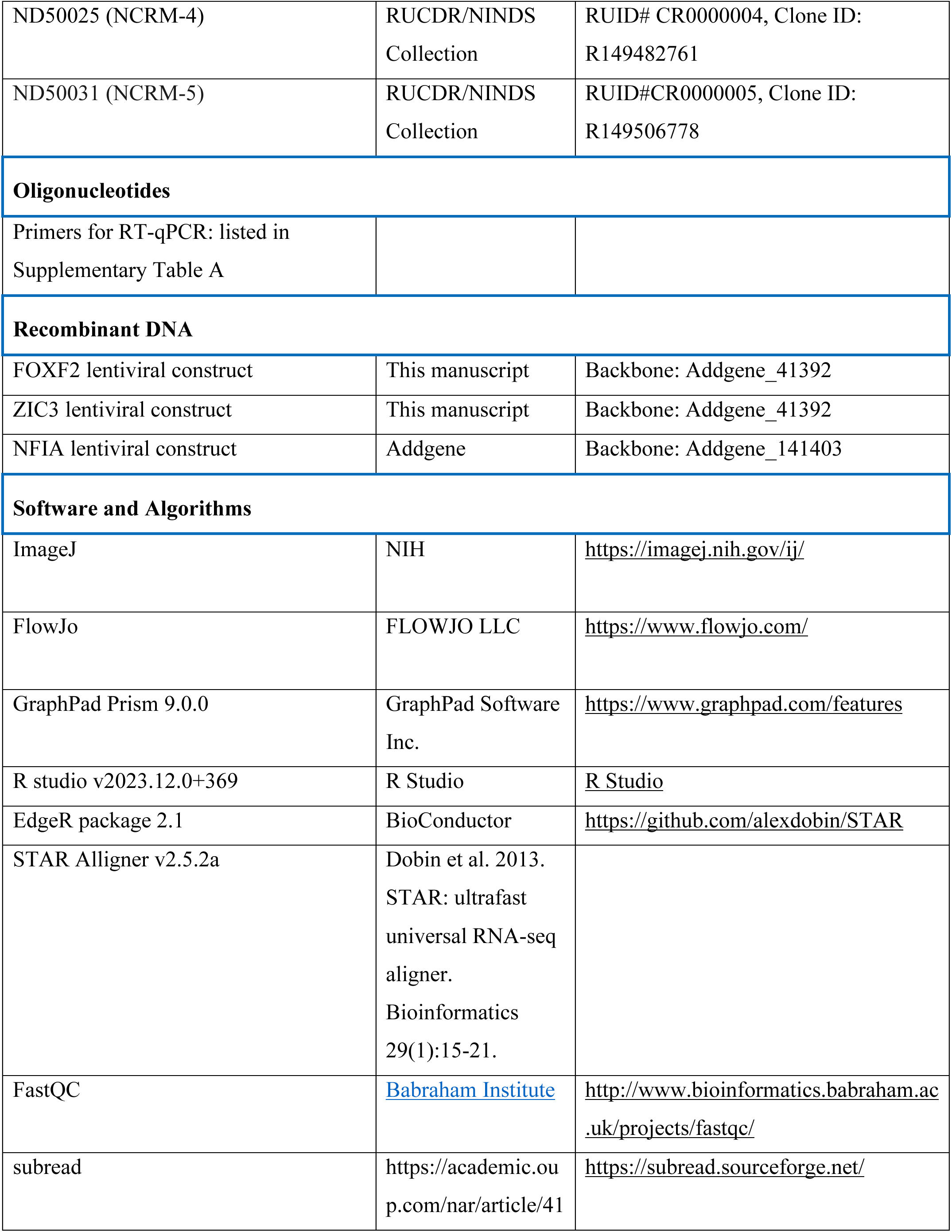
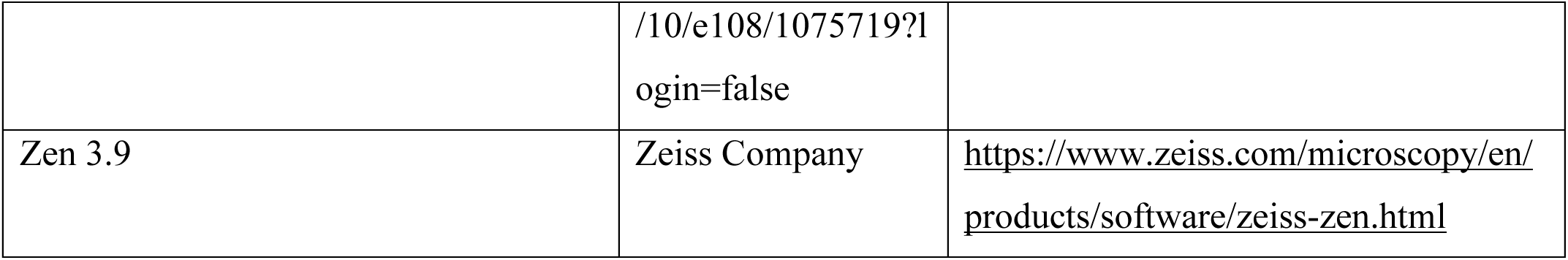

### Resource availability

For further information and requests for resources, reagents, or code, please contact corresponding authors Andrew Sproul and Dritan Agalliu.

### Materials availability

Plasmids from this study are available upon reasonable request. Please refer to STAR*Methods section for resources.

### Experimental Models

#### Human iPSC line maintenance

For all primary experiments, human control iPSC line IMR90-cl.4 was used (WiCell)^70^. For additional experiments, human control iPSC line FA0000010 (FA10), ND50031 (NCRM-5), and ND50025 (NCRM-4) were used (RUCDR). IMR90-cl.4 and NCRM-4 were derived from female donors, while FA10 and NCRM-5 were from male donors. For differentiation into bpECs or NPCs, iPSCs were maintained on feeder culture of irradiated CF1 mouse embryonic fibroblasts (MEFs) with Human Embryonic Stem cell Medium (HUESM): knockout DMEM, 20% knockout serum replacement (Thermo Fisher), 2mM Glutamax (Thermo Fisher), 0.1 mM MEM non-essential amino acids (Thermo Fisher), 1% penicillin-streptomycin (Thermo Fisher), and 0.1 mM β-mercaptoethanol (Thermo Fisher) supplemented with 20 ng/mL bFGF (R&D) and passaged when 80-90% confluent. Feeder-free hiPSCs were maintained in StemFlex medium (Thermo Fisher) containing penicillin-streptomycin (Thermo Fisher) on Cultrex-coated plates (Bio-Techne). Cultures were passaged using ReLeSR (StemCell Technologies) every 3-4 days.

#### Directed differentiation of hiPSCs into BBB-primed endothelial cells (bpECs)

hiPSCs were differentiated into bpECs (our term) as previously reported with slight modifications^35^. hiPSCs were dissociated with collagenase-IV, and cell aggregates were plated in chemically defined media (CDM): 50% IMDM (Thermo Fisher), 50% Ham’s F12 (Thermo Fisher), 5 mg/mL bovine serum albumin (Thermo Fisher), 15 ug/mL transferrin (Sigma-Aldrich), 0.1 mM β-mercaptoethanol (Thermo Fisher), 7 ug/mL insulin (Sigma-Aldrich), 1% penicillin-streptomycin (Thermo Fisher) on 1 ug/mL fibronectin (Sigma-Aldrich) coated 10 cm dishes from one well of a 6-well plate to one 10 cm dish. CDM was supplemented with 10 ng/mL BMP4 (R&D) for 1.5 days to induce mesodermal differentiation and 50 ng/mL BMP4 and 20 ng/mL bFGF (R&D) for the next 3.5 days for further expansion of mesodermal lineage cells. From days 5 to 10, endothelial differentiation was initiated by supplementing CDM media with 50 ng/mL VEGF (R&D) and 100 ng/mL thymosin-β4. Endothelial cells were purified at day 10-11 by FACS for CD31^+^/CDH5^+^ cells. These endothelial cells, now termed bpECs, were plated on collagen IV/fibronectin coated plates at a density of 30,000 to 35,000 cells/cm^2^ and fed EC medium (Human Endothelial Serum Free Medium (Thermo Fisher) supplemented with 1% human serum (Sigma-Aldrich) supplemented with 50 ng/mL VEGF165 (R&D), 20 ng/mL bFGF (R&D), and 10 μM SB431542 (Sigma-Aldrich). bpECs were expanded and passaged 1:2 to 1:3 every 7-10 days and generally used for five passages.

#### Reprogramming of bpECs into rBMECs

bpECs were plated at 30,000-35,000 cells/cm^2^ on collagen IV/fibronectin coated plates using EC media supplemented with 50 ng/mL VEGF165, 20 ng/mL bFGF and 10 uM SB431542. On day 2, bpECs were transduced with FOXF2 (~1×10^6^ lentiviral p24 units/mL of medium) and ZIC3 (~7.5×10^5^ lentiviral p24 units/mL of medium) as determined by Lenti-X GoStix (Takara). Successfully infected cells were selected with 1 ug/mL puromycin (Sigma) for 24 to 48 hours. Infected bpECs were purified via FACS for CD31^+^/VE-cadherin^+^ cells on day 14. FACS-purified and infected bpECs, referred to as rBMECs, were plated on collagen IV/fibronectin coated plates 25,000-35,000 cells/cm^2^ and expanded until confluent. rBMECs were passaged at high density using 1:2 splits every 7-10 days and used within five passages.

#### Generation of lentiviral vectors and lentiviruses

FOXF2 and ZIC3 were subcloned into pLex_307, which was a gift from David Root (Addgene), and verified by Sanger sequencing (Azenta). FUW-tetO-NFIA was a gift from Lorenz Studer (Addgene)^71^. FUdeltaGW-rtTA was a gift from Konrad Hochedlinger (Addgene)^72^. psPAX2 was a gift from Didier Trono (Addgene). VSV.G was a gift from Tannishtha Reya (Addgene)^73^. Lentiviral particles for FOXF2, ZIC3, NFIA, and rtTA were generated in HEK293 as described previously^74^, using the psPAX2/VSV.G 2^nd^ generation system. Efficacy of lentiviral particles was confirmed by infection and subsequent qPCR.

#### Fluorescence-activated cell sorting (FACS)

Cells were dissociated into single cell suspension using Accutase (ThermoFisher) and washed with base medium (CDM for EC differentiation at day 10 and EC media for rBMECs). Cell suspensions were incubated with FACS buffer (10% KOSR in DPBS) with fluorescently tagged antibodies (CD31-FITC, VE-cadherin-APC) for 30 min at room temperature protected from light. Cell suspensions were washed with FACS buffer and cell pellets were resuspended for FACS by filtering through 0.45 μM filters (Fisher Scientific) to remove cell aggregates. bpECs or rBMECs were sorted on the Sony MA900 cell sorter and double positive CD31^+^/VE-cadherin^+^ cells were subsequently expanded.

#### Primary endothelial cell, pericyte, and astrocyte cultures

HBMECs (Cell Systems) were cultured in Endothelial Cell Growth Medium MV 2 (Promocell). PCs were cultured in Pericyte Media (PM, ScienCell), and ACs were cultured in Astrocyte Media (AM, ScienCell). HBMECs and PCs were plated onto collagen IV and fibronectin coated plates, while ACs were plated onto either Matrigel or Cultrex-coated plates. All cells were supplemented with media every other day and passaged using Accutase once 90% confluent.

#### Generation of iPSC-derived pericytes

Briefly, feeder-free iPSCs were differentiated into neural crest pericytes following an established protocol, where iPSCs were treated with a neural crest differentiation media supplemented with CHIR99021 and subsequently treated with Pericyte Media (ScienCell)^54^. Cells were later sorted on CD146+/PDGFRβ+ to ensure a homogeneous population prior to introduction into the BBB triculture system.

#### Generation of iPSC-derived astrocytes

hiPSCs were differentiated into a neural progenitor cells (NPCs) using dual-SMAD inhibition in embryoid bodies as previously described^56^. On day −1, NPCs were dissociated using Accutase and plated at a density of 15,000 cells/cm^2^ in N2/B27 media on Cultrex-coated plates. On day 0, the media was replaced with supplemented Astrocyte Media (2% fetal bovine serum, and proprietary astrocyte growth factor supplement) (ScienCell). On day 1, differentiating cells were transfected with lentiviral particles carrying dox-inducible NFIA and rtTA (Addgene). Cells were fed every other day using astrocyte media supplemented with doxycycline (2μg/mL) (Sigma-Aldrich). Cells were split at 90-95% confluence roughly once every seven days and re-plated at a density of 15,000 cells/cm^2^ for 30 days. At this stage, astrocytes were split 1:3 once 90-95% confluent and were expanded, banked, and utilized.

#### RNA isolation and RT-qPCR

RNA was extracted using the RNeasy Plus Micro Kit (Qiagen) and resuspended in 30 µl of RNAse-free water. RNA quantity and quality were assessed with a Nanodrop One (ThermoScientific). For qPCR, 100-250 ng of RNA was transcribed into cDNA using the Maxima First Strand cDNA Synthesis Kit (Thermo Fisher). Gene expression was assessed using the FAST SYBR Green Master Mix (ThermoFisher Scientific) with primer sets (Integrated DNA Technologies) for target genes on a QuantStudio 7 Flex system (Applied Biosystems) and CFX Opus 384 (Bio-Rad). Gene expression for target genes were normalized to GAPDH and fold changes were calculated as ΔΔC_t_.

#### Western blotting

Endothelial cells were harvested using RIPA Lysis Buffer supplemented with Halt protease and phosphatase inhibitors single-use cocktails (Thermo Fisher) on ice. 10 μg of lysates and 1x Laemmli sample buffer/5% β-mercaptoethanol were boiled at 95° C for 5 min. SDS-PAGE was performed by loading samples on Bolt 4–12% gradient Bis-Tris gels using Bolt MOPS buffer (Thermo Fisher). Nitrocellulose membranes were first blocked with Superblock TBS buffer/0.1% Tween 20 Thermo Fisher) for 1h, and then incubated overnight at 4° C with the following primary antibodies diluted in Superblock/0.1% Tween 20: Claudin-5 (1:1000; Thermo Fisher), Occludin (1:1000; Thermo Fisher), VCAM-1 (1:1000, Abcam), pSTAT3 (1:1000, BioLegend), STAT3 (1:1000, BioLegend) and GAPDH (1:5000; Thermo Fisher). After 3 washes with TBS/0.1% Tween-20, membranes were then incubated with the appropriate HRP-conjugated secondary antibody (1:5000, Thermo Fisher) for 1 hour, and washed 3 more times with TBS/0.1% Tween-20. Pierce ECL western blotting substrate or SuperSignal West Pico PLUS chemiluminescent substrate (Thermo Fisher) were used to develop chemiluminescence and blots were imaged using the Bio-Rad Chemidoc MP system.

#### Transendothelial electrical resistance (TEER) measurement

TEER measurements were carried out using the ECIS Z-theta instrument on the 96W20idf array at 4000 Hz frequency. Briefly, 20,000 cells were plated on poly-D-lysine and collagen IV/fibronectin-coated 96-well electrode array plates in the respective medium for each cell type. Resistance was recorded for 250-300 hours and medium was changed every other day. The resistance curve was generated using PRISM (GraphPad) and the area under the curve (AUC) was measured either for last 100 hours of recording or for the duration of oAβ42 or cytokine treatment (72hr or 24hr) from 6-8 wells using the AUC PRISM function.

#### Tracer permeability assay

Tracer permeability assay for determination of paracellular permeability was performed as previously described^75^. rBMECs, bpECs and hBMECs were cultured on 12-well polycarbonate Transwell inserts (0.4 uM, Corning) for 10 days to ensure a complete monolayer. Growth factors were removed for 24 hours prior to assay. A full media change was performed on bottom chamber (1.5 mL) and all 3 tracers (NaF-10 ug/mL, 3 kDa dextran - 25 ug/mL and 70 kDa dextran-25 ug/mL) were added on top chamber in 0.5 mL medium to start the assay. As soon as tracers were added, 100 uL medium was collected from bottom chamber as 0 min collection and replaced with 100 uL fresh media for each well, followed by serial collection for every 10 min for up to 60 min. Fluorescent intensities were measured and clearance slopes were calculated using:

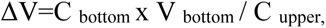

C_bottom and_ C_upper_ are fluorescent intensities measured for each tracer in top and bottom chamber respectively, V_bottom_ is volume of media in bottom chamber (1.5 mL)

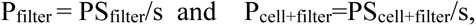

P_filter_ and P_cell+filter_ are permeability coefficient for blank insert and insert with cells, PS_filter_ and PS_cell+filter_ are which can be obtained by regression plot of ΔV against time for blank insert and insert with cells.

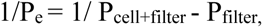

efficient for endothelial cell monolayer

#### Wound scratch assay

Cells were plated at a density of 100K/well (24-well plate). Once confluent, cells were fed their standard media absent VEGF and bFGF supplementation. Monolayers were scratched in the center of each well using the Autoscratch automated scratch machine (BioTek). Wells were then treated with either control media, 100 ng/mL VEGF, or an angiogenic cocktail consisting of VEGF (37.5 ng/ml), bFGF (37.5 ng/ml), and HGF (37.5 ng/ml). Plates were cultured and imaged automatically every 6 hours using the Cytation5 plate reader (BioTek) and software, for HBMECs (24 hrs) and bpECs and rBMECs (36 hrs) during culture in a BioSpa incubator (BioTek). Each well was quantified for % surface area of the wound utilizing an ImageJ plug-in for identifying the area of the scratch^76^, which was then normalized to the original size of the wound for each well. To compare cell types, we calculated an angiogenic potential, where:

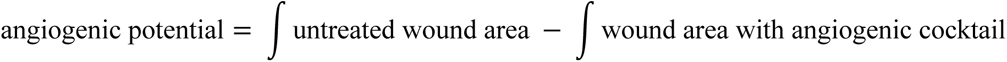

#### Ki67 quantification assay

After the wound scratch assay was performed, cells were fixed with 4% PFA. Cells were blocked as described above and stained with Ki67 antibody (1:500). Cells were labeled with a GFP secondary antibody and DAPI. Wells were imaged for DAPI and GFP fluorescence on the BioSpa as done previously. Each well was quantified for proliferative index, where:

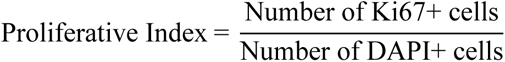

Cells were identified as Ki67 or DAPI positive using the annotation machine learning tool from the QuPath software^77^.

#### Uptake and transporter assays

As previously described^60^, cells were plated at a density of 100K/well (24-well plate) and subcultured until confluent. First, cells incubated with either 1 mg/mL unlabeled BSA or 1 mg/mL BSA-Alexa 594 for 1 hour at 37° C. Cells were then dissociated into single cell suspension, stained for eFluor 780, fixed with 4% PFA, and analyzed for Alexa594 median fluorescence intensity (MFI). For the transporter assay, cells were incubated with Rhodamine 123 (10 uM) for 1 hr at 37° C. To assess the effect of P-glycoprotein (Pgp) efflux, cells were pretreated with PSC-833 (a specific Pgp inhibitor) for 1 hr prior to Rhodamine treatment. Cells were dissociated into single cell suspension, stained for eFluor 780, fixed with 4% PFA, and analyzed for Rhodamine MFI. NPCs were utilized as a positive control due to their low level of Pgp efflux. Data were normalized to the unstained MFI for each cell type, which we termed fold change in MFI. To calculate Pgp efflux, we calculated the percent change in MFI with and without PSC-833:

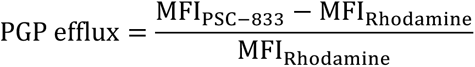

#### Amyloid-β42 oligomerization and treatment

Amyloid-β42 was monomerized, stored and later oligomerized as previously described^78,79^. In brief, Aβ(1-42) (rPeptide, 1 mg) vial was allowed to equilibrate to room temperature. Peptides were resupended in ice-cold hexafluoroisopropanol (Sigma-Aldrich) to achieve 1 mM concentration directly in a vacuum-sealed vial using a Hamilton syringe. The suspension was divided into 8 equal aliquots by volume using a Hamilton syringe into low protein binding Eppendorf tubes, incubated at room temperature for 2 hours and dried on speedvac at 800g for 25°C. Upon drying monomerized Aβ(1-42) films can be stored at −80°C in sealed tubes. For generating oligomers, Eppendorf tubes containing Aβ(1-42) peptide films were equilibrated to room temperature, resuspended in DMSO to achieve 5 mM concentration and sonicated at 17°C for 10 min for complete resuspension. Oligomerization was induced by adding phenol-red free F12 to achieve final concentration of 100 uM and incubated at 4°C for 24 to 48 hours. rBMEC monolayers were treated with oAβ at 500 nM or 1 uM concentrations for the indicted period of time and F12 was used a vehicle control.

#### 3D microfluidic BBB culture model

OrganoPlate 3-lane-64 plates (MIMETAS) were used to generate a 3D microfluidic system of the BBB. In each chip, the center channel was filled with an ECM gel, an endothelial tubule was generated in the right channel, and a mixture of iPSC-derived astrocytes and pericytes was seeded in the left channel and allowed to migrate towards the endothelial cells during culture upon a bidirectional OrganoFlow rocker (MIMETAS).

##### ECM generation and endothelial channel coating

On day −1, the center ECM channel was filled with a 4:1 ratio of collagen I (5 mg/ml) and collagen IV solution (5 mg/mL), incubated at 37° C for 15 min, and covered with PBS to prevent collagen gel from drying. The right channel inlet was coated using fibronectin (10 ug/mL) and collagen IV (10 ug/mL) overnight.

##### Endothelial cell seeding

On day 0, endothelial cells were singularized and resuspended to achieve a 15,000 cells/uL concentration. Cells were seeded in the right channel outlet at a density of 30,000 cells/chip. Plate was placed on a 75° plate stand from the manufacturer and incubated at 37° C for four to five hours to allow endothelial cells to settle against the gel. Then, outlets were replenished with endothelial media and the plate was placed on the OrganoFlow rocker to alternate between −7 to +7° every 8 minutes. After a day or two, endothelial cells formed an open tubule in the right channel.

##### Astrocyte and pericyte channel coating and cell seeding

On day 1, the left channel inlet was coated with Matrigel. Primary astrocytes (ACs) and iPSC-derived ACs were singularized and resuspended in AM-PM media (2:1 ratio of AM:PM) to achieve a 15,000 cells/uL concentration. Primary pericytes (PCs) and iPSC-derived iPCs were similarly singularized and resuspended in AM-PM to achieve a 7,500 cells/uL concentration. Primary astrocytes and primary pericytes were mixed at a ratio of 1:1 and seeded in the left channel outlet for each chip to achieve 15,000 ACs and 7,500 PCs per chip. The plate was rested on the stand for four to five hours at 37° C to allow pericytes and astrocytes to settle against the gel. Finally, outlets were replenished with AM-PM and the plate was placed back on the OrganoFlow rocker for continuous culture until Day 10. Media was replenished every other day. Beginning on day 6, chips were replenished with their standard EC media without bFGF and VEGF supplementation.

#### Tracer permeability assay

On day 10 post seeding, a 70-kDa dextran (0.5 ug/mL) and Biocytin tracer (0.5 ug/mL) were introduced into the right inlet and outlet wells (40 uL and 30 uL, respectively). The gel inlet, left inlet, and left outlet wells were filled with 20 uL media without tracer. Then, the plate was placed onto the OrganoFlow rocker and imaged every 15 minutes in the LI-COR. The fluorescence intensity for each well was quantified with the Odyssey SA system. Analysis was performed as the normalized fluorescence intensity over the ECM channel center well, which represented the leakage from the right vascular channel into the left brain channel:

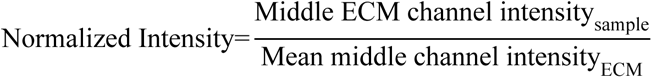

where ECM chips only had a control gel, representing the leakage without cells present. Samples were analyzed using a two-way ANOVA test.

#### 3D TEER assay

Chips were assayed for TEER on day 10 post seeding. After the plate was introduced into the OrganoTEER device, TEER values were measured for the vascular channel using the “low TEER” setting, which is suitable for endothelial cells, per the manufacturer. Measurements for each condition were compared using a one-way ANOVA test.

### Immunofluorescence and microscopy

#### Staining of monolayer endothelial cells, pericytes, and astrocytes

For tight junction staining, endothelial cells were fixed with ice cold ethanol for 30 minutes and acetone for 1 minute. For all other staining, ECs, PCs, and ACs were fixed in 4% PFA for 15 min. Fixed monolayers were then washed twice in PBS, blocked for an hour in blocking buffer (5% goat or donkey serum, 0.3% Triton X-100 in PBS), and incubated overnight with appropriate primary antibodies in a diluting solution at 4° C (5% goat or donkey serum; 0.01% Triton X-100 in 1 x PBS). The following day, cells were stained with secondary antibodies in 1 x PBS for 1 hour at room temperature and DAPI (1:5000) for 10 minutes. Imaging was performed on a ZEISS LSM700 confocal microscope using 10x (Plan-Apochromat 10x objective lens) and 20x (Plan Apochromat 20x objective lens).

#### Immunostaining of 3D MIMETAS cultures

On day 10 post seeding, MIMETAS chips with fixed with either 4% PFA for 15 minutes for standard staining or ethanol/acetone for tight junction visualization. For fixation and blocking, solution was introduced as follows: 100 uL in right channel inlet, 50 uL in right channel outlet, 50 uL in left channel inlet, 50 uL in left channel outlet. Plate was placed on the OrganoFlow for rocking during staining. Chips were blocked for an hour in blocking buffer, and incubated overnight with primary antibodies in diluting solution at 4° C as follows: 25 uL in right channel inlet and outlet; 15 uL in left channel inlets and outlets. Cells were washed and incubated with secondary antibodies and DAPI as above. Then, chips were post-fixed with 4% PFA for 15 minutes prior to microscopic imaging.

### Bulk RNA sequencing, alignment and analysis

Total RNA was extracted as previously described. RNA samples underwent quality assessment using the Qubit 2.0 Fluorometer, and RNA integrity was assessed using the Agilent TapeStation 4200. Subsequently, RNA sequencing libraries were prepared using the NEBNext Ultra II RNA Library Prep Kit following the manufacturer’s protocol. The samples were sequenced in a 2×150bp Paired-End (PE) configuration using the Illumina NovaSeq instrument.

Previously published samples were obtained from Gene Expression Omnibus (GEO) datasets, including GSE138025, GSE40291, GSE57662, GSE131039, GSE97575, GSE73721, GSE97324, GSE138025, GSE122588, GSE129290, GSE97100, GSE126449, GSE108012, GSE151976, GSE85403, GSE82207, GSE77250, GSE82207, GSE157852, GSE243193, GSE225549, and GSE227501. Quality control for the samples was conducted using FastQC, followed by adapter trimming with Skewer (v0.2.2). The resulting reads were aligned to the human reference genome (hg38) using the STAR aligner, and counts were generated using subread.

Normalization was performed using the trimmed mean of M-values (TMM) method implemented in the EdgeR R package. Fragments per kilobase of transcript per million mapped reads (FPKM) and Log2 counts per million (cpm) matrices were generated to account for library size variations. Differential expression analysis was conducted considering a p-adjusted value < 0.05 and a log2fold change less than −0.25 or greater than 0.25.

Principal component analysis and Pearson correlation values were computed using the resulting Log_2_ cpm values. Heatmaps were generated using Min-Max scaling on normalized counts.

