## Supplemental Figures S1-S4 for "Generation of hiPSC-derived brain microvascular endothelial cells using a combination of directed differentiation and transcriptional reprogramming strategies"

### SUPPLEMENTARY INFORMATION

### SUPPLEMENTARY FIGURES AND LEGENDS

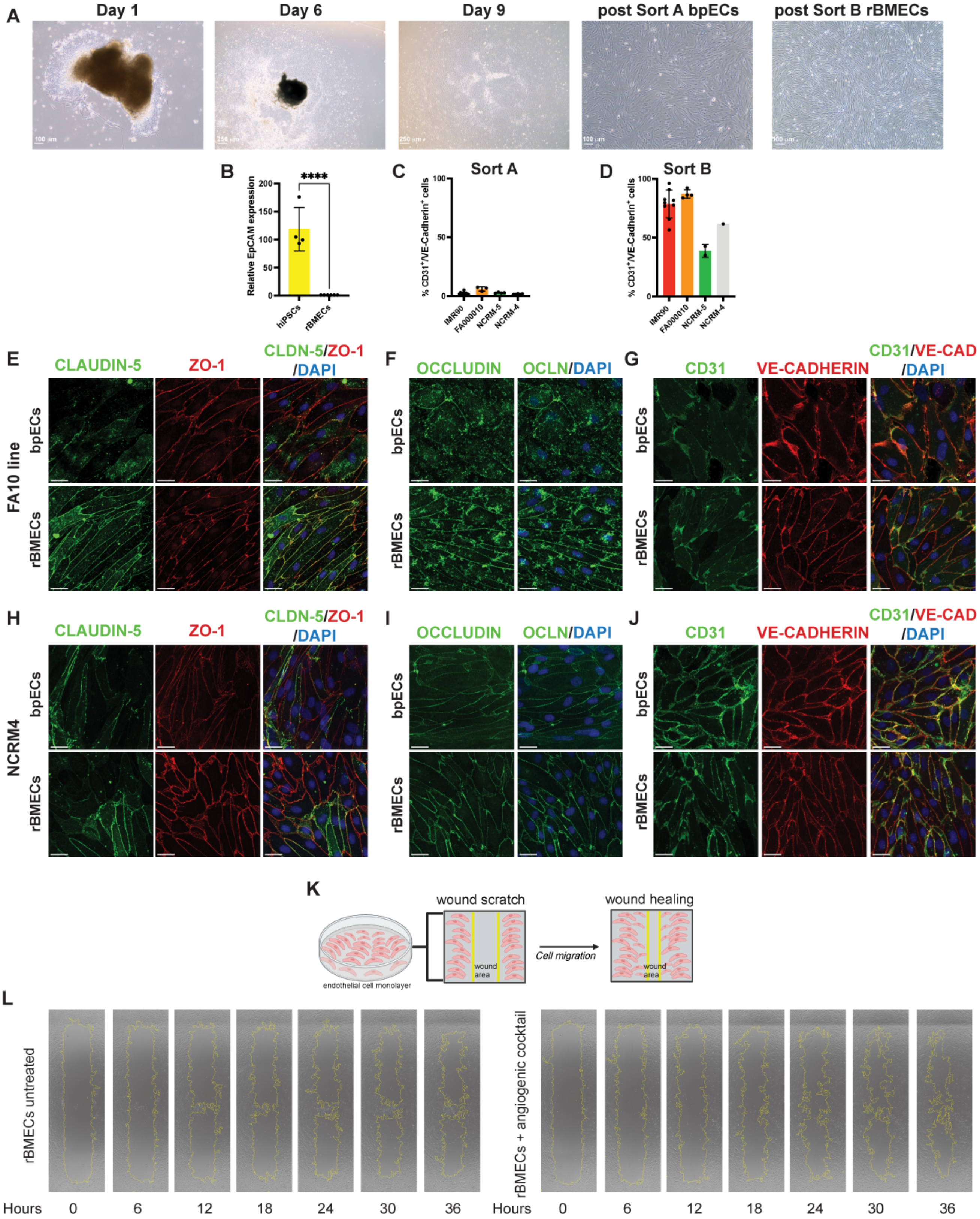

**Figure S1 (related to Figure 1). rBMECs generated from multiple iPSC lines express BBB-tight junction proteins. A)** Representative brightfield images show human induced pluripotent stem cells (hiPSC) on select days (D1, D6, and D9) during differentiation and bpECs (post-sort A), and rBMECs (post-sort B). **B)** Dotted bar graph of the relative EpCAM surface expression in hiPSCs and rBMECs. Mean ± SD, n = 4 from 2 independent experiments, unpaired Student t-test, ****p<0.0001. **C-D)** Dotted bar graphs of the quantification of CD31$^{+}$/VE-cadherin$^{+}$ cells obtained from additional hiPSC lines for bpECs [post sort A **(C)**] and rBMECs [post-sort B **(D)**]. Each dot represents an independent differentiation. **E-G)** Representative immunofluorescence (IF) images of bpECs and rBMECs derived from the FA10 hiPSC line for **(E)** Claudin-5 (green) and ZO-1 (red), **(F)** Occludin (green), **(G)** CD31 (green) and VE-Cadherin (red). DAPI (blue) labels nuclei in all merged images. Scale bar = 25 µm. **H-J)** Representative IF images of bpECs and rBMECs derived from the NCRM-4 hiPSC line for **(H)** Claudin-5 (green) and ZO-1 (red), **(I)** Occludin (green), **(J)** CD31 (green) and VE-Cadherin (red). DAPI (blue) labels nuclei in all merged images. Scale bar = 25 µm. **K)** Schematic of the wound scratch assay. **L)** Representative images show rBMEC wound scratch healing over a 36-hour time course in the absence (left) or presence (right) of the angiogenic cocktail (see Methods for details).

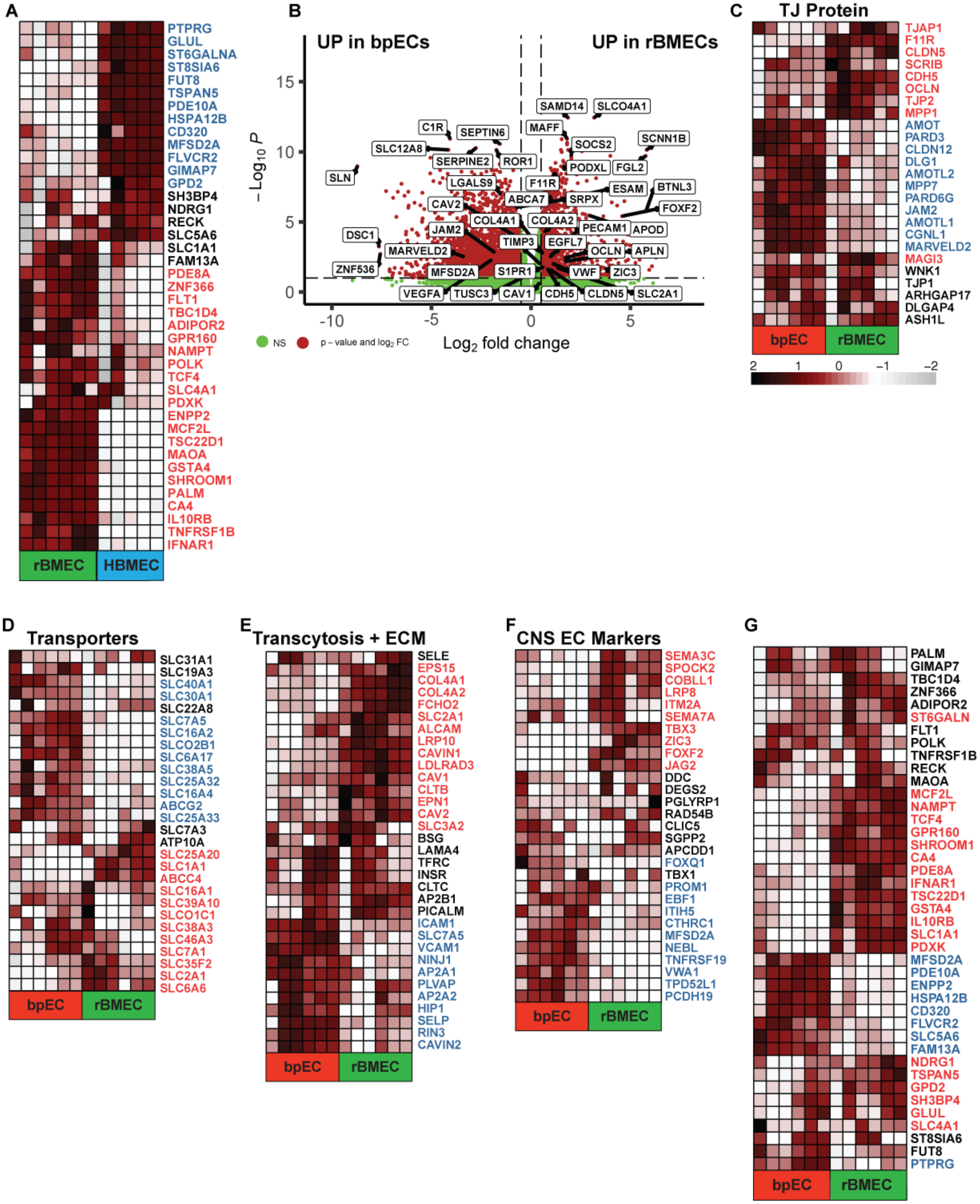
A
PTPRG
GLUL
ST6GALNA
ST8SIA6
FUT8
TSPAN5
PDE10A
HSPA12B
CD320
MFSD2A
FLVCR2
GIMAP7
GPD2
SH3BP4
NDRG1
RECK
SLC5A6
SLC1A1
FAM13A
PDE8A
ZNF366
FLT1
TBC1D4
ADIPOR2
GPR160
NAMPT
POLK
TCF4
SLC4A1
PDXK
ENPP2
MCF2L
TSC22D1
MAOA
GSTA4
SHROOM1
PALM
CA4
IL10RB
TNFRSF1B
IFNAR1
rBMEC
HBMEC
B
UP in bpECs
UP in rBMECs
15
10
5
0
−Log10 P
-10
-5
0
5
Log2 fold change
NS
p − value and log2 FC
SAMD14
SLCO4A1
C1R
SEPTIN6
MAFF
SCNN1B
SLC12A8
SOCS2
SERPINE2
ROR1
PODXL
FGL2
SLN
F11R
ESAM
BTNL3
LGALS9
ABCA7
SRPX
FOXF2
CAV2
COL4A1
COL4A2
PECAM1
APOD
JAM2
DSC1
TIMP3
EGFL7
OCLN
APLN
MARVELD2
S1PR1
VWF
ZIC3
ZNF536
MFSD2A
VEGFA
TUSC3
CAV1
CDH5
CLDN5
SLC2A1
C
TJ Protein
TJAP1
F11R
CLDN5
SCRIB
CDH5
OCLN
TJP2
MPP1
AMOT
PARD3
CLDN12
DLG1
AMOTL2
MPP7
PARD6G
JAM2
AMOTL1
CGNL1
MARVELD2
MAGI3
WNK1
TJP1
ARHGAP17
DLGAP4
ASH1L
bpEC
rBMEC
2
1
0
−1
−2
D
Transporters
SLC31A1
SLC19A3
SLC40A1
SLC30A1
SLC22A8
SLC7A5
SLC16A2
SLCO2B1
SLC6A17
SLC38A5
SLC25A32
SLC16A4
ABCG2
SLC25A33
SLC7A3
ATP10A
SLC25A20
SLC1A1
ABCC4
SLC16A1
SLC39A10
SLCO1C1
SLC38A3
SLC46A3
SLC7A1
SLC35F2
SLC2A1
SLC6A6
bpEC
rBMEC
E
Transcytosis + ECM
SELE
EPS15
COL4A1
COL4A2
FCHO2
SLC2A1
ALCAM
LRP10
CAVIN1
LDLRAD3
CAV1
CLTB
EPN1
CAV2
SLC3A2
BSG
LAMA4
TFRC
INSR
CLTC
AP2B1
PICALM
ICAM1
SLC7A5
VCAM1
NINJ1
AP2A1
PLVAP
AP2A2
HIP1
SELP
RIN3
CAVIN2
bpEC
rBMEC
F
CNS EC Markers
SEMA3C
SPOCK2
COBLL1
LRP8
ITM2A
SEMA7A
TBX3
ZIC3
FOXF2
JAG2
DDC
DEGS2
PGLYRP1
RAD54B
CLIC5
SGPP2
APCDD1
FOXQ1
TBX1
PROM1
EBF1
ITIH5
CTHRC1
MFSD2A
NEBL
TNFRSF19
VWA1
TPD52L1
PCDH19
bpEC
rBMEC
G
PALM
GIMAP7
TBC1D4
ZNF366
ADIPOR2
ST6GALN
FLT1
POLK
TNFRSF1B
RECK
MAOA
MCF2L
NAMPT
TCF4
GPR160
SHROOM1
CA4
PDE8A
IFNAR1
TSC22D1
GSTA4
IL10RB
SLC1A1
PDXK
MFSD2A
PDE10A
ENPP2
HSPA12B
CD320
FLVCR2
SLC5A6
FAM13A
NDRG1
TSPAN5
GPD2
SH3BP4
GLUL
SLC4A1
ST8SIA6
FUT8
PTPRG
bpEC
rBMEC

**Figure S2 (related to Figure 2). The rBMEC transcriptome shows higher expression of key BBB identity genes compared to the bpEC transcriptome. A)** Heatmap of the relative expression of the core BBB genes between rBMEC and primary HBMECs. The upregulated genes in rBMECs are shown in red and the upregulated genes in HBMECs are shown in blue. **B)** Volcano plot for differentially expressed genes in rBMECs compared to bpECs. All BBB-specific genes listed are statistically significant ($p < 0.05$, $|logFC| > 0.25$). **C-F)** Heatmap of the relative expression of BBB-specific genes related to **(C)** TJ proteins, **(D)** transporter genes, **(E)** transcytosis and ECM-related genes, and **(G)** CNS endothelial cell marker genes in bpECs and rBMECs. **G)** Heatmap of the relative expression of the core BBB genes between bpECs and rBMECs. The upregulated genes in rBMECs are in red, and the upregulated genes in bpECs are in blue. The scale for all heatmaps is shown in panel **C**.

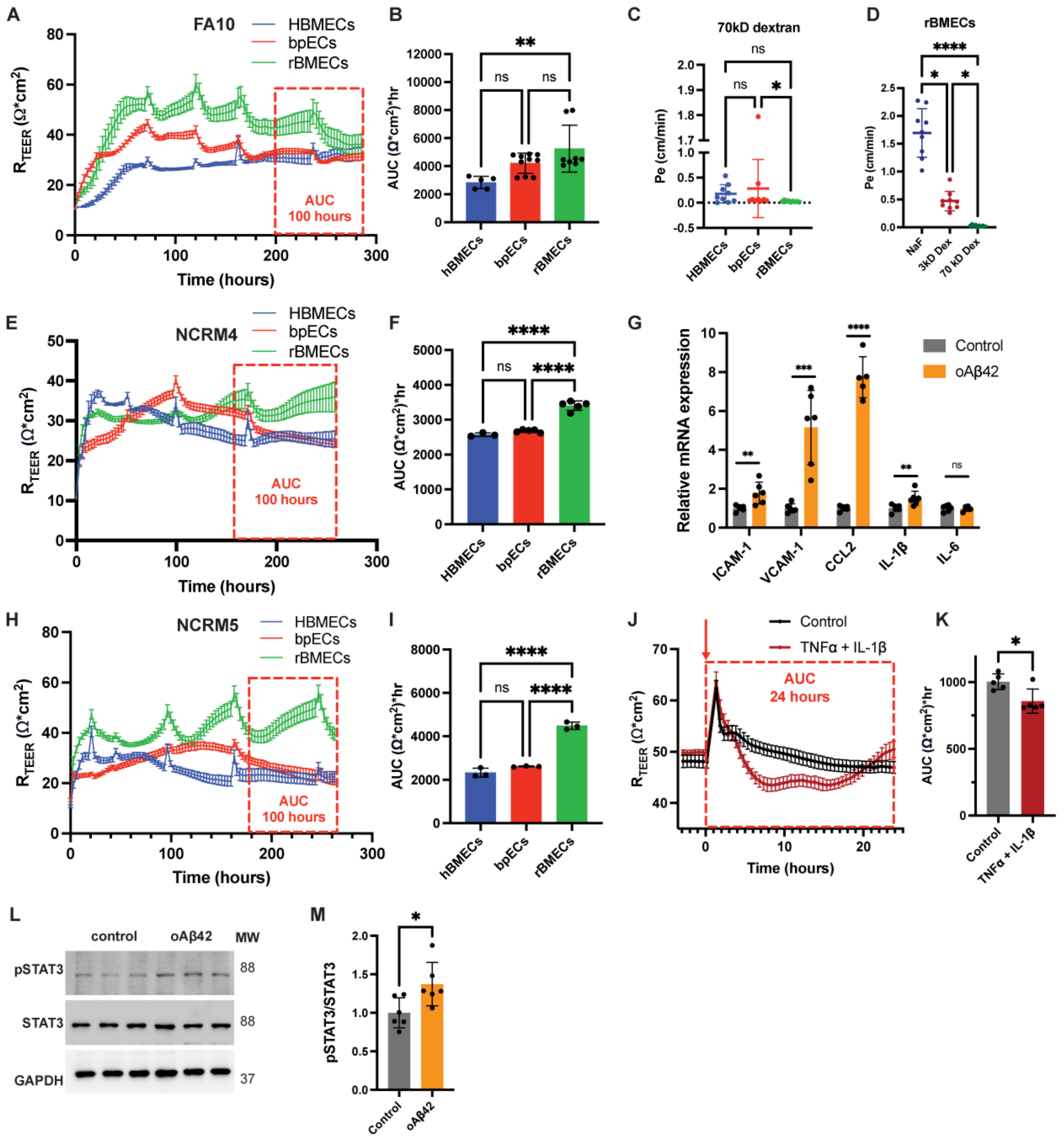

A
FA10
HBMECs
bpECs
rBMECs
AUC
100 hours
Time (hours)
B
C
70kD dextran
D
rBMECs
NaF
3kD Dex
70 kD Dex
E
NCRM4
F
G
Relative mRNA expression
Control
oAβ42
ICAM-1
VCAM-1
CCL2
IL-1β
IL-6
H
NCRM5
I
J
Control
TNFα + IL-1β
AUC
24 hours
K
L
control
oAβ42
MW
pSTAT3
STAT3
GAPDH
M
pSTAT3/STAT3

**Figure S3 (related to Figure 3). rBMECs can be derived from multiple iPSC-lines with improved barrier properties and respond robustly to an inflammatory stimulus. A-B)** Representative ECIS TEER recording and dotted bar graph of the area under the curve (AUC) quantification for FA10 hiPSC-derived bpECs, rBMECs, and primary HBMECs. The red box indicates the period used for the AUC measurements. Mean ± SD, n =5-10 from 3 experiments, one-way ANOVA, $^{**}p<0.01$. **C)** Dotted bar graph of the transwell permeability to 70 kDa dextran (Pe). Mean ± SD, n = 9 from 3 independent experiments, ANOVA with Kruskal-Wallis Test, $^{*}p<0.05$. **D)** Permeability (Pe) comparison of sodium fluorescein (NaF), 3 kDa dextran permeability, and 70 kDa dextran across monolayers of rBMECs. Mean ± SD, n = 9 from 3 independent experiments, ANOVA with Kruskal-Wallis Test, $^{*}p<0.05$, $^{****}p<0.0001$. **E-F)** Representative ECIS TEER recording and bar graph of the AUC quantification for NCRM-4 hiPSC-derived bpECs, rBMECs, and primary HBMECs. Mean ± SD, n = 3-5 from 1 experiment, one-way ANOVA, $^{****}p<0.0001$. **G)** Bar graph of the relative expression of select inflammatory genes in rBMECs by RT-qPCR after 72 hr treatment with 1 uM oAβ42. Mean ± SD, n = 6 for 2 experiments, unpaired Student t-test, $^{**}p<0.01$, $^{***}p<0.001$, $^{****}p<0.0001$. **H-I)** Representative ECIS TEER recording and bar graph of the AUC quantification for NCRM-5 hiPSC-derived bpECs, rBMECs, and HBMECs. Mean ± SD, n = 3 from 1 experiment, one-way ANOVA, $^{****}p<0.0001$. **J-K)** Representative ECIS TEER recording and bar graph of the AUC quantification for rBMECs treated with recombinant TNF-α (10 ng/mL) and IL-1β (10 ng/mL). Mean ± SD, n = 5 from 2 experiments, unpaired t-test, $^{*}p<0.05$. **L-M)** Representative western blots and quantification of phospho-STAT3 and total-STAT3 in rBMECs after a 6-hour treatment with 500 nM oAβ42. Mean ± SD, n = 6 from 2 experiments, unpaired t-test, $^{*}p<0.05$.

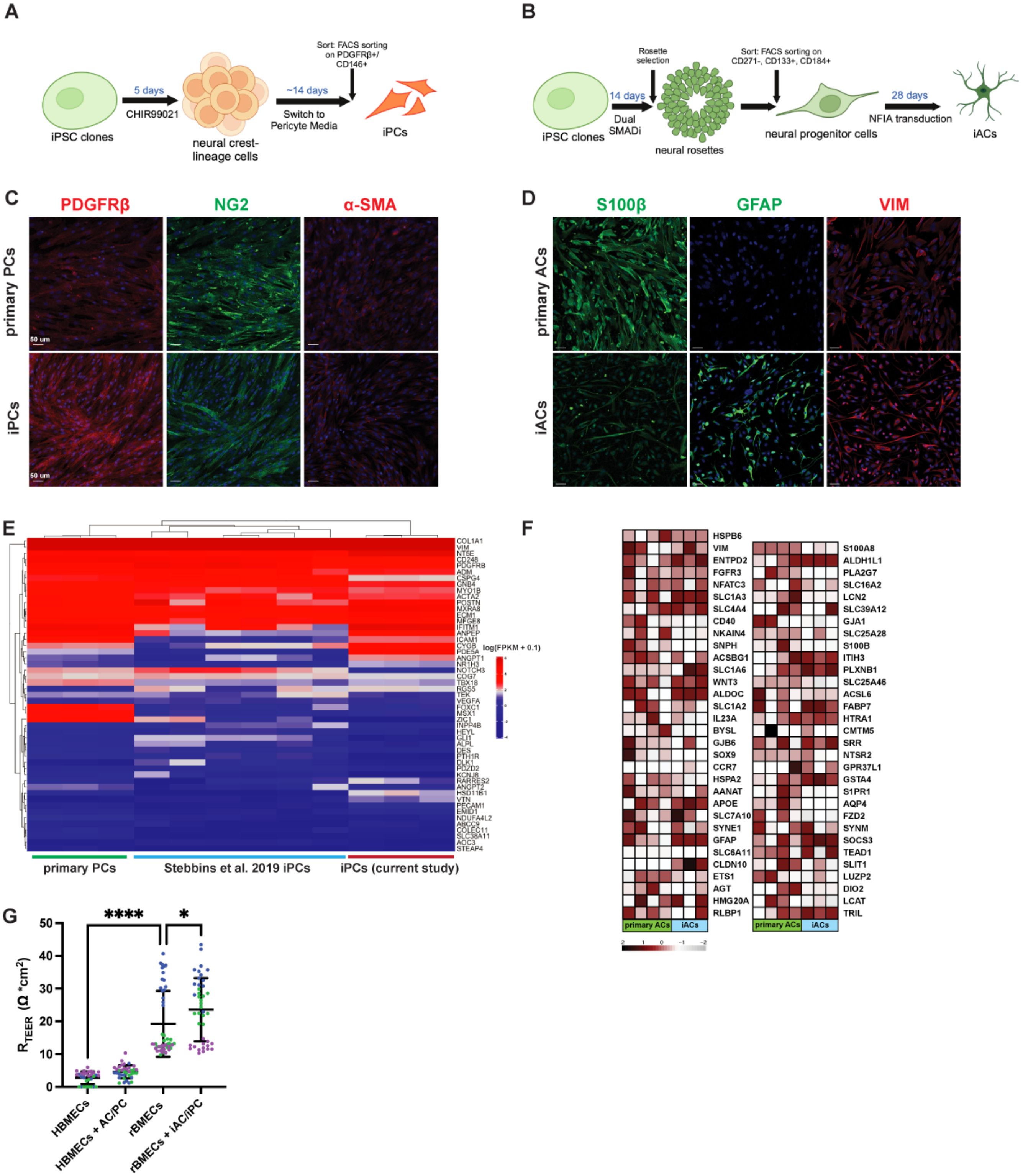
A
5 days
CHIR99021
~14 days
Switch to Pericyte Media
Sort: FACS sorting on PDGFRβ+/ CD146+
iPSC clones
neural crest-lineage cells
iPCs
B
Rosette selection
14 days
Dual SMADi
Sort: FACS sorting on CD271-, CD133+, CD184+
28 days
NFIA transduction
iPSC clones
neural rosettes
neural progenitor cells
iACs
C
PDGFRβ
NG2
α-SMA
primary PCs
iPCs
50 um
D
S100β
GFAP
VIM
primary ACs
iACs
E
log(FPKM + 0.1)
primary PCs
Stebbins et al. 2019 iPCs
iPCs (current study)
F
HSPB6
VIM
ENTPD2
FGFR3
NFATC3
SLC1A3
SLC4A4
CD40
NKAIN4
SNPH
ACSBG1
SLC1A6
WNT3
ALDOC
SLC1A2
IL23A
BYSL
GJB6
SOX9
CCR7
HSPA2
AANAT
APOE
SLC7A10
SYNE1
GFAP
SLC6A11
CLDN10
ETS1
AGT
HMG20A
RLBP1
S100A8
ALDH1L1
PLA2G7
SLC16A2
LCN2
SLC39A12
GJA1
SLC25A28
S100B
ITIH3
PLXNB1
SLC25A46
ACSL6
FABP7
HTRA1
CMTM5
SRR
NTSR2
GPR37L1
GSTA4
S1PR1
AQP4
FZD2
SYNM
SOCS3
TEAD1
SLIT1
LUZP2
DIO2
LCAT
TRIL
primary ACs
iACs
G
****
*
R_TEER (Ω *cm²)
50
40
30
20
10
0
HBMECs
HBMECs + AC/PC
rBMECs
rBMECs + iAC/iPC

**Figure S4 (related to Figure 4). Characterization of hiPSC-derived pericytes and astrocytes. A, B)** Schematic diagrams for generation of hiPSC-derived pericytes and hiPSC-derived astrocytes. **C)** Representative immunofluorescence staining of primary and hiPSC-derived pericytes (iPCs) for PDGFRβ (red), NG2 (green), and α-SMA (green). **D)** Representative IF images of hiPSC-derived astrocytes (iACs) for S100β (green), GFAP (green) and Vimentin (red). **E)** Heatmap showing relative mRNA expression of pericyte markers from primary pericytes, hiPSC-derived pericytes generated in this study, and hiPSC-derived pericytes generated from a previously published dataset (Stebbins et al., 2019). **F)** Heat map showing mRNA expression of astrocyte markers from primary astrocytes, hiPSC-derived astrocytes generated in this study, and hiPSC-derived astrocytes generated from other groups. Data is presented as log(z-score). **G)** 3D TEER measurement for mono-and co-cultured EC tubules. Dots are color coded by experiment to show experiment-to-experiment variation in TEER values; n = 30-48 chips over three experiments, one-way ANOVA, ****<0.0001, *<0.05.

### SUPPLEMENTARY TABLES AND LEGENDS

**Table S1 (related to Figure 2 and Figure S2). A)** PCA components for all cell types. **B)** PCA components for endothelial cells only. **C)** FPKM values for all bulk RNA sequencing samples. **D)** Differentially expressed genes between rBMECs and HBMECs. **E)** Differentially expressed genes between rBMECs and bpECs.

| Gene | Forward Primer | Reverse primer |
|---|---|---|
| ICAM-1 | CAATGTGCTATTCAAACTGCCC | CAGCGTAGGGTAAGGTTCTTG |
| VCAM-1 | TCTACGCTGACAATGAATCCTG | AGGGCCACTCAAATGAATCTC |
| E-Selectin | AAGTTCGCCTGTCCTGAAG | CAGAAAGTCCAGCTACCAAGG |
| CCL2 | CCTCCAGCATGAAAGTCTCTG | TCTGCACTGAGATCTTCCTATTG |
| TNFα | ACTTTGGAGTGATCGGCC | GCTTGAGGGTTTGCTACAAC |
| IL-1β | ATGCACCTGTACGATCACTG | ACAAAGGACATGGAGAACACC |
| IL-6 | CAACCTGAACCTTCCAAAGAT | ACCTCAAACTCCAAAAGACCAG |
| GAPDH | TGAAGGTCGGAGTCAACGGATTTGG | CATGTAGGCCATGAGGTCCACCAC |

**Table S2 (related to STAR methods).** The list of primers used for the RT-PCR experiments described in Figures 3L and S3G.

### SUPPLEMENTARY MOVIES

**Supplemental Movie 1.** 3D projection of rBMECs in MIMETAS labeled with VE-cadherin (green) and CD31 (red), associated with Figure 4C.

**Supplemental Movie 2.** 3D projection of rBMECs in MIMETAS labeled with ZO-1 (green) and Claudin-5 (red), associated with Figure 4C.

**Supplemental Movie 3.** 3D NVU showing iPCs (PDGFRβ) and iACs (S100β) migrating from the brain channel towards the blood channel, associated with Figure 4D.

**Supplemental Movie 4.** 3D projection of rBMECs, iPCs, and iACs in MIMETAS labeled with stained with ZO-1 (red), S100b (green), and PDGFRb (blue), associated with Figure 4E.
